# Endothelial cell-derived, secreted long non-coding RNAs *Gadlor1* and *Gadlor2* aggravate pathological cardiac remodeling via intercellular crosstalk

**DOI:** 10.1101/2022.09.19.508486

**Authors:** Merve Keles, Steve Grein, Natali Froese, Dagmar Wirth, Felix A. Trogisch, Rhys Wardman, Shruthi Hemanna, Nina Weinzierl, Philipp-Sebastian Koch, Stefanie Uhlig, Santosh Lomada, Gesine M. Dittrich, Malgorzata Szaroszyk, Ricarda Haustein, Jan Hegermann, Abel Martin-Garrido, Johann Bauersachs, Derk Frank, Norbert Frey, Karen Bieback, Julio Cordero, Gergana Dobreva, Thomas Wieland, Joerg Heineke

**Author notes:** Address for Correspondence: Prof. Dr. Joerg Heineke, European Center for Angioscience (ECAS), Department of Cardiovascular Physiology, Medical Faculty Mannheim, Heidelberg University, Ludolf-Krehl-Straße 7 – 11, 68167 Mannheim, Germany. National Heart and Lung Institute, Faculty of Medicine, Imperial College London, London, United Kingdom (Merve Keles). Department of Pharmacology and Toxicology, University Medical Center Göttingen, Göttingen, Germany (Gesine M. Dittrich). These authors contributed equally (Merve Keles, Steve Grein).

## Abstract

**Background:** Pathological overload triggers maladaptive myocardial remodeling that leads to heart failure. Recent studies have shown that long non-coding RNAs (lncRNAs) regulate cardiac remodeling. This study investigates two recently discovered, secreted lncRNAs, *Gadlor1* and *Gadlor2* (*Gadlor 1/2*).

**Methods:** We generated compound *Gadlor*1/2 knock-out (KO) mice and compared their response to pressure overload by transverse aortic constriction (TAC) to that of wild-type (WT) littermates. Endothelial cells, fibroblasts and cardiomyocytes were isolated from the hearts of both genotypes after TAC and their transcriptome was investigated by RNA sequencing. *Gadlor* target proteins were identified by RNA antisense purification coupled with mass spectrometry (RAP-MS) in cardiomyocytes. In addition, we investigated the effects of cardiac overexpression of *Gadlor1*/*2*.

**Results:** *Gadlor1/2* are jointly upregulated in failing mouse hearts as well as in the myocardium of heart failure patients. Cardiac overexpression of *Gadlor1* and *Gadlor2* aggravated myocardial dysfunction and enhanced hypertrophic and fibrotic remodeling in mice exposed to pressure overload. Compound *Gadlor1/2* KO mice, in turn, exerted markedly reduced myocardial hypertrophy, fibrosis and dysfunction, but more angiogenesis during short and long-standing pressure overload. Paradoxically, *Gadlor1/2* KO mice suffered from sudden death during prolonged overload, possibly due to cardiac arrhythmia. *Gadlor1* and *Gadlor2*, which are mainly expressed in endothelial cells (ECs) in the heart, where they inhibit pro-angiogenic gene-expression, are strongly secreted within extracellular vesicles (EVs). These EVs transfer *Gadlor* lncRNAs to cardiomyocytes, where they bind and activate calmodulin-dependent kinase II, induce pro-hypertrophic gene-expression and enhance calcium re-uptake into the sarcoplasmic reticulum.

**Conclusion:** *Gadlor1* and *Gadlor2* are lncRNAs that are mainly enriched in EC-derived EVs and are jointly upregulated in mouse and human hearts during pathological overload. We reveal a crucial endothelial cell-cardiomyocyte crosstalk, which aims at restoring calcium homeostasis in cardiomyocytes during overload at the cost of aggravated hypertrophy and fibrosis.

## Introduction

Despite recent advances in therapeutic options, chronic heart failure is still one of the leading causes of global mortality, in part due to insufficient means to counteract maladaptive cardiac remodeling.^1–3^ Cardiac remodeling comprises global changes in heart shape, reduced function and arrhythmia that are the consequence of cardiomyocyte hypertrophy, altered angiogenesis and myocardial fibrosis.^4, 5,6, 7^ The majority of current therapeutic options target protein-coding genes; however, recent studies revealed the significance of non-coding transcripts in cardiovascular development and disease conditions.^5, 8, 9^

Long non-coding RNAs (lncRNAs) are a diverse group of non-coding transcripts that are longer than 200 nucleotides, and that in general do not encode for proteins.^10^ Due to their complex structure, lncRNAs can act, for instance, as chromatin modifiers, affect protein functions, or alter mRNA stability or processing.^5^ Thereby lncRNAs typically affect gene expression, alternative splicing or signaling pathways.^11, 12^ Recent studies have identified several lncRNAs with crucial roles in cardiac remodeling and heart failure.^11, 13–16^ Since non-coding transcripts are detectable in human body fluids such as in serum, circulating microRNAs (miRNAs) and lncRNAs are currently being investigated as clinical biomarkers, as well as for being paracrine mediators of cellular communication during stress conditions.^17–19^ Intra-myocardial communication is a highly regulated process governing the complex physiology of the heart, whereby proper synchronization is required, for example, between endothelial cells (ECs) and cardiomyocytes, which rely on EC lined capillaries to provide nutrients and oxygen, but also paracrine factors regulating cardiomyocyte growth and function^20–22^. Cellular communication during pathological remodeling can be mediated by proteins, but also by the secretion and transfer of nanoscale extracellular vesicles (microvesicles and exosomes, termed small extracellular vesicles, EVs, in here) that carry non-coding RNAs and alter the behavior of recipient cells. ^23–28,29,30^ It is currently emerging that EVs transport lncRNAs between cells and thereby affect endothelial cell function, cardiac fibrosis and hypertrophy, but more work is needed.^26, 28, 31^

Here, we studied two related and secreted lncRNAs *Gadlor1* and *Gadlor2* (GATA-downregulated long non-coding RNA), which are jointly upregulated after endothelial deletion of GATA2.^32^ We found that the expression of both *Gadlor* lncRNAs are significantly increased in hearts of mice after transverse aortic constriction (TAC), but also in the myocardium and serum of patients suffering from chronic heart failure. Moreover, our results revealed that *Gadlor1* and *Gadlor2* are secreted within EVs predominantly by cardiac ECs, from which they are mainly taken up by cardiomyocytes, although they also have autocrine/intracrine effects on ECs. In cardiomyocytes, *Gadlor1/2* impact calcium handling, hypertrophy and gene expression, while triggering anti-angiogenic effects in ECs. Our study reveals that secreted *Gadlor* lncRNAs aggravate cardiac remodeling but might protect from arrhythmias by enhancing calcium cycling.

## Methods

Detailed descriptions of materials and methods are provided in the **Supplemental Material**.

### Human Samples

Studies on human heart tissue samples were approved by the Institutional Ethical Board of Massachusetts General Hospital (US), where samples were collected from patients with end-stage heart failure undergoing cardiac transplantation.^33^ Control heart tissue samples were obtained from healthy organ volunteers when the organ was not eligible for transplantation, or from the victims of traffic accidents. Human serum samples were obtained from healthy blood donors, or from aortic stenosis patients before replacement of the aortic valve (clinical data are available in **Supplemental Table 1**), where all donors were provided a written informed consent for the collection and use of samples, that had received approval from the Institutional Review Board of Christian-Albrechts-Universität Kiel (File number: A174/09).

### Animal Models and Studies

All studies including the use and care of animals were performed with the permission of the Regional Council Karlsruhe and the Lower Saxony State Office for Consumer Protection and Food Safety, Germany, with approved protocols 35-9185.81/G-144/18, I-22/03, 33.9-42502-12-10/0016, 33.19-42502-04-14/1403 and 33.8-42502-04-16/2356. Female and male animals were used in similar proportions throughout the study, except for the experiments in Figure 3, where only male mice were used. Systemic *Gadlor* knock-out mice (*Gadlor*-KO) were generated by deletion of the respective region of mouse chromosome 16 using a CRISPR-Cas9-based strategy to achieve homozygous deletion. Pressure overload was induced in 8-10 weeks old mice by transverse aortic constriction (TAC) and maintained for 2 weeks for short-term and 8-12 weeks for long-term studies. Mice were monitored with echocardiography before, and during pressure overload studies for assessment of cardiac function.

### Primary Cell Isolation

Isolation of endothelial cells and fibroblasts from adult mouse hearts was performed with MACS-magnetic beads from Miltenyi Biotec based on the manufacturer’s protocol. Briefly, the tissue lysate was incubated with CD146 microbeads (Miltenyi Biotec, 130-092-007) before being passed through magnetic columns (Miltenyi Biotec, MS columns 130-042-201) for positive selection of EC. The flow through was incubated with feeder removal microbeads (Miltenyi Biotech, 130-095-531) and eluted after multiple washing steps to collect fibroblasts (FB). Isolation of adult cardiac myocytes was achieved by using a Langendorff perfusion system. Isolated cells were stored as frozen cell pellets for later RNA or protein isolation for corresponding experiments.

### Extracellular Vesicle Isolation

Isolation of EVs was performed with ultracentrifugation from the supernatant of C166 cells that were cultured with EV-depleted FCS. Initially, cell debris and large vesicles (apoptotic bodies) were removed by centrifugations at 300xg and 10.000xg, respectively. Subsequently, small EVs (including microvesicles and exosomes) were collected with 100.000xg ultracentrifugation for 90 minutes and similarly washed with PBS, before a second round of centrifugation was performed. For details on EV characterization and visualization, see **Supplemental Material.**

### Sarcomere Contractility and Calcium Transient Measurements

Isolated adult cardiomyocytes were plated on laminin coated (10 µg/cm2) 35 mm dishes (MatTek 10 mm Glass bottom dishes, P35G-1.5-10-C) and gently washed 1 hour later with MEM medium without butanedione monoxime (BDM). Contractility and calcium transient measurements were performed by using Ion-Optix Multicell High-throughput (HT) System chamber, where the cells were simultaneously paced (18 V, 2.5 Hz, 4 ms impulse duration), and assessed for sarcomere length and contraction. Calcium handling was recorded after incubation with 1 µM fura-2, AM (Invitrogen, F1221).

### RNA antisense purification coupled with mass spectrometry (RAP-MS)

To identify the protein interaction partners of *Gadlor* lncRNAs we performed RAP-MS. The 5’ biotinylated antisense *Gadlor1* and *Gadlor2* probes were pooled in the experiment and the sequences are listed in **Supplemental Table 5**. HL-1 cardiomyocytes overexpressing *Gadlor1* and *Gadlor2* were used, where each experimental group contained three replicates without cross-linking (as a negative control to identify non-specific ‘background’ proteins) and four replicates with UV cross-linked condition by using 150 mJ/cm^2^ of 254 nm UV light. After hybridization of biotin labelled probes, lysates were cleared with streptavidin coated magnetic beads. The captured protein samples were identified by TMT labelling followed by liquid chromatography-mass spectroscopy (LC-MS/MS) by the EMBL Proteomics core facility.

### RNA sequencing and Bioinformatics

RNA isolation was performed from isolated cells as described at indicated time points (2 weeks after TAC or sham surgery). Quality control of RNA samples (Agilent 2100 Fragment Analyzer), and library preparation (DNBSEQ Eukaryotic Strand-specific mRNA library) were performed by BGI, Hong Kong. Bulk RNA sequencing from different cardiac cells was performed as stranded and single-end with 50 base sequence read length by BGI with Illumina HiSeq 2500. For details of RNAseq analysis, see **Supplemental Material.**

### Data Availability

The authors declare that the data supporting the findings of this study are available within the paper and its supplementary information. RNA sequencing data sets were deposited in National Center for Biotechnology Information’s (NCBI) Gene Expression Omnibus (GEO) repository with the accession number GSE213612.

### Statistical Analysis

Data are shown as mean ± standard error of the mean (SEM). Data analysis and statistical analysis were performed with GraphPad Prism software (Version 8). An unpaired 2-tailed Student t-test was used for comparing 2 groups. Comparing multiple groups for one or multiple conditions was performed with one-way ANOVA and two-way ANOVA, respectively, and Fisher’s LSD post-hoc test was conducted when applicable. For non-parametric data sets, the Mann-Whitney and Kruskal-Wallis tests were used to compare two groups and multiple groups, respectively. Values of p<0.05 were considered statistically significant.

## Results

### Increased expression of *Gadlor* lncRNAs in cardiac pressure overload in mice and human failing hearts

Earlier observations by our group had identified two previously unknown lncRNAs *Gadlor1* and *Gadlor2*, which were markedly and jointly upregulated in cardiac ECs of endothelial-specific GATA2 knock-out mice.^32^ *Gadlor1* (AK037972) and *Gadlor2* (AK038629) are located in close proximity to each other on mouse chromosome 16, and are embedded after exon1 of the *Lsamp* (limbic system-associated membrane protein) gene in its intronic region **(Figure 1A)**. *Lsamp* is mainly expressed in the brain, and we did not detect significant *Lsamp* expression in the heart or cardiac ECs, and *Lsamp* expression patterns did not change in response to *Gadlor* knock-out in multiple organs **(Figure S1A-C)**. *Gadlor1* and *Gadlor2* are not matched with any known mouse peptides (BlastP, The National Center for Biotechnology Information (NCBI) database), and we verified that they are non-coding transcripts by using the CPAT algorithm to assess protein-coding probability **(Figure S1D)**. Cardiac *Gadlor1* and *2* expression is low in the direct postnatal period, but both transcripts become upregulated in parallel in the adult (2 months old) heart **(Figure 1B)**. Cellular expression analysis among the main cell types in the heart revealed the highest *Gadlor1* and *Gadlor2* expression in ECs, followed by fibroblasts (FBs), while the lowest levels were detected in cardiomyocytes (CMs) **(Figure 1C)**. In these analyses, *Gadlor 1* and *2* lncRNA levels were also significantly enriched in ECs compared to whole heart tissue.

**Figure 1:**
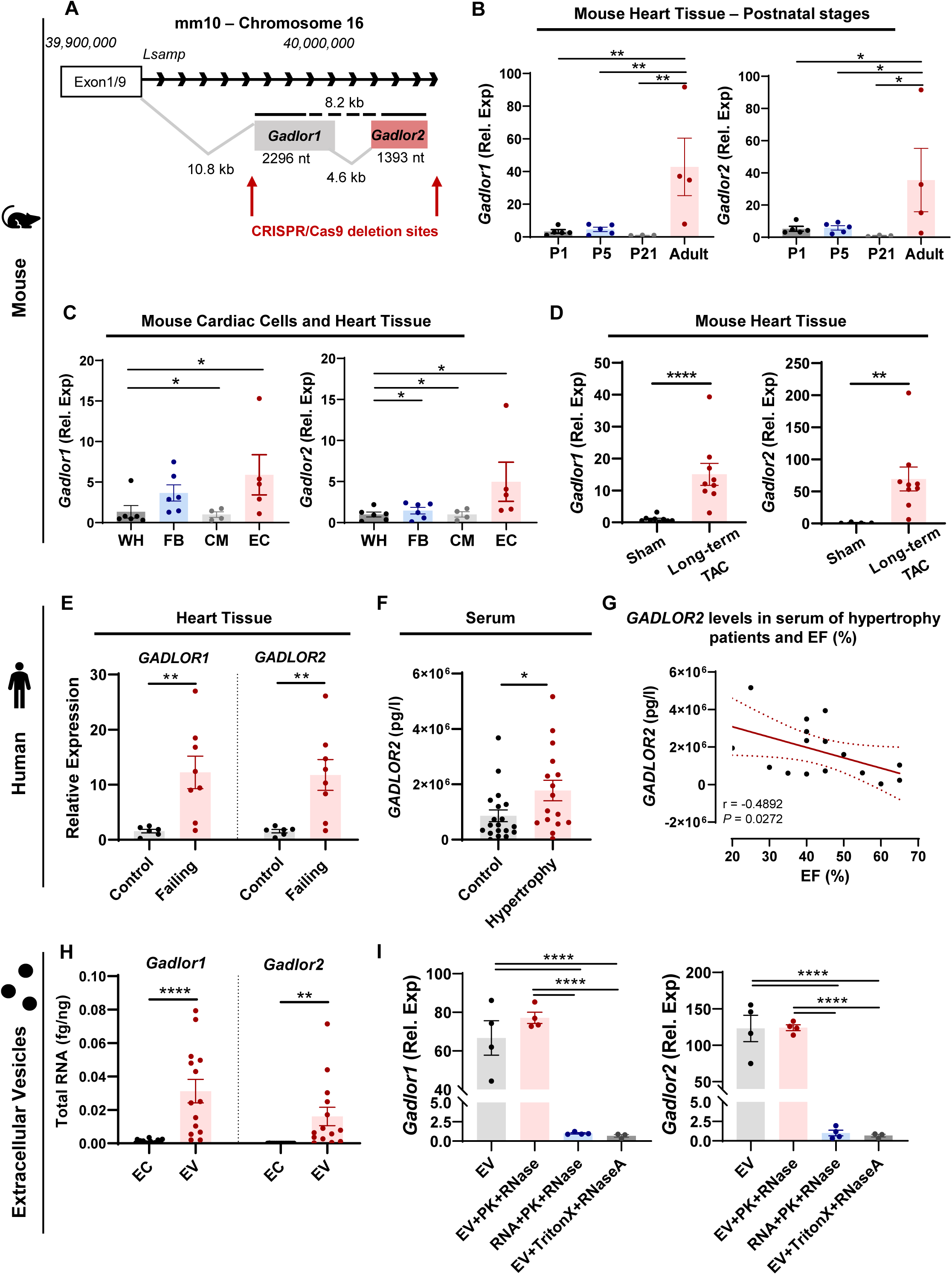
*Gadlor* lncRNAs *are* mainly expressed in cardiac endothelial cells (ECs) and their expression is highly increased in mice after pressure-overload and in human failing hearts. **A.** Representative scheme of AK037972 *(Gadlor1)* and AK038629 *(Gadlor2)* locus on mouse chromosome 16 and positions of the CRISPR/ Cas9 deletion sites used to generate *Gadlor-*KO mouse line. **B.** *Gadlor1* and *Gadlor2* expression levels in mouse cardiac tissue from different post-natal developmental stages (P1 – n=5, P5 – n= 5, P21 – n=3 and adult (9-weeks old) – n=4), and in **C.** different cardiac cell-types (EC: endothelial cells – n=5, FB: fibroblast – n=6, CM: cardiomyocytes – n=4) and heart tissue (WH: whole heart – n=6). **D.** *Gadlor* lncRNA levels in mouse cardiac tissue after long-term (12 weeks, n=9) TAC (transverse aortic constriction) compared to sham. **E.** *GADLOR1* and *2* expression in control (healthy, n=6) and human failing heart tissue (n=8) samples that were obtained from aortic stenosis patients. **F.** Detection of *GADLOR2* levels (pg/l: picogram/liter) in human serum of healthy volunteers (n=19) and patients with aortic stenosis (n=16). **G.** Correlation analysis of *GADLOR2* levels (pg/l) with ejection fraction (%) of hypertrophy patients (n=16), (Pearson correlation, Pearson r= −4892 and p-value = 0.0272). **H.** Absolute quantification of total RNA amount of *Gadlor1* and *Gadlor2* in ECs and EC-derived EVs (fg/ng: femtogram/ nanogram). **I.** Detection of *Gadlor* lncRNAs in EC-derived EVs with proteinase K (PK), RNase or Triton-X as indicated. Data are shown as mean±SEM. Data normality was evaluated with the Shapiro-Wilk test. *P*-values were calculated with Student’s t-test for parametric (or Mann-Whitney for non-parametric) for comparing two groups, and one-way ANOVA was applied for comparing multiple groups followed with Fisher’s LSD post-hoc test. *p-value<0.05, **p-value<0.01, ***p-value<0.001, ****p-value<0.0001.

Next, we assessed *Gadlor1/2* RNA levels in cardiac disease. Myocardial *Gadlor1* and *2* expression was markedly and jointly upregulated in the chronic phase (12 weeks) after TAC versus sham **(Figure 1D)**. Predominant endothelial expression and the increased abundance during pressure overload were confirmed by fluorescent in situ hybridization **(Figure S1E-F)**.

The conservation of *Gadlor1/2* RNA between human, mouse and rat was analyzed with the EMBOSS-Water platform **(Figure S2)**, revealing high conservation between the three species. Importantly, *GADLOR1/2* expression was strongly induced in the myocardium of patients suffering from advanced heart failure **(Figure 1E)**. We established an assay to specifically detect and quantify *GADLOR2* concentrations in human serum, which were significantly elevated in patients with heart failure due to aortic stenosis **(Figure 1F)**. In this patient cohort, high *GADLOR2* serum concentrations were found mainly in patients with low left ventricular ejection fraction, and a negative correlation between these two parameters was identified **(Figure 1G).** We were not able to establish a similar assay to quantify *GADLOR1* in human serum. Taken together, our results demonstrate *Gadlor1* and *Gadlor2* as lncRNAs that are co-upregulated in diseased mouse and human hearts.

### *Gadlor* lncRNAs are enriched in EC-derived extracellular vesicles (EVs)

Since *GADLOR2* was detectable in the serum of heart failure patients, and secreted EVs can contain lncRNAs,^26, 29^ we hypothesized that *Gadlor*1/2 might be secreted within EVs. To test our hypothesis, the abundance of *Gadlor1*/*2* was analyzed in cultured primary mouse cardiac ECs and EVs collected and purified from the supernatant of the ECs by ultracentrifugation **(Figure S3A)**. Initially, we characterized the isolated EVs with electron microscopy (EM), nanoparticle tracking analysis (NTA) and FACS. We found typical microvesicles and exosomes in EM, and the average size of the particles was slightly larger than 100 nm in diameter by NTA, which showed that they were in the range of microvesicles and exosomes (together called EVs in this study) **(Figure S3B-C)**. Flow cytometry assessment also confirmed the presence of CD63, CD9 and CD54 as common cell surface markers for EVs **(Figure S3D)**.

Interestingly, the absolute concentration of *Gadlor1/2* was markedly higher in cardiac EC-derived EVs compared to cardiac EC themselves **(Figure 1H)**. EV *Gadlor1* and *2* RNA levels were not affected by proteinase K (PK), or by RNase A, which indicated *Gadlor1/2* were encapsulated within vesicles and protected from enzyme degradation **(Figure 1I)**. Indeed, *Gadlor* lncRNAs were not protected upon direct incubation with PK and RNase A, or when EVs were dissolved by Triton-X, before RNase A was applied **(Figure 1I)**. In addition to primary cardiac ECs, also C166 mouse ECs secreted *Gadlor* RNAs in EVs **(Figure S3E)**. Altogether, these results suggest that *Gadlor1* and *Gadlor2* are highly enriched and encapsulated in EC derived EVs.

### *Gadlor*-KO mice are protected from cardiac hypertrophy and dysfunction during pressure-overload

To investigate the role of *Gadlor* lncRNAs, systemic *Gadlor* knock-out mice (*Gadlor*-KO) were generated by deletion of the respective region of mouse chromosome 16 (including *Gadlor* 1 and 2, since both were consistently jointly regulated in our analyses) by using a CRISPR-Cas9-based strategy **(Figure 1A**, **Figure 2A)**. Unchallenged *Gadlor*-KO mice were indistinguishable from their wild-type littermates (WT) based on appearance, body weight, heart weight, and baseline echocardiographic analysis **(Figure S4A-D)**. We detected, however, a mild increase in liver and decrease in brain weight in KO animals compared to WT littermates (**Figure S4E**).

**Figure 2:**
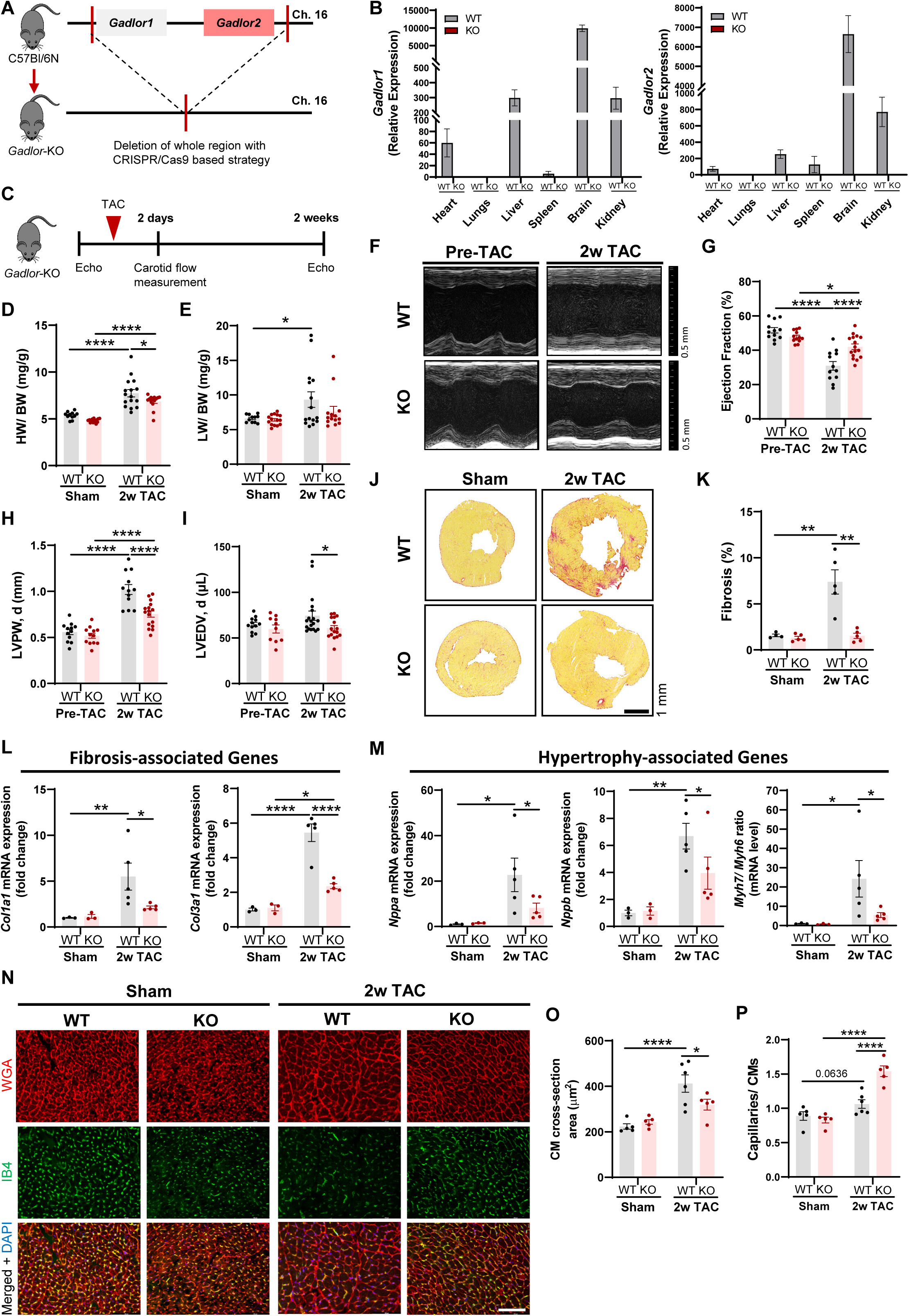
Deletion of *Gadlor* lncRNAs reduces cardiac hypertrophy and fibrosis after cardiac pressure overload. **A.** Scheme of *Gadlor-*KO mouse line generation by systemic deletion of the region on chromosome 16 containing both *Gadlor1* and *Gadlor2* by a CRISPR/Cas9 approach. **B.** Validation of *Gadlor-*KO mouse line by assessing the relative expression of *Gadlor*s in indicated organs in adult WT and *Gadlor-*KO mice (n≥8). The difference in *Gadlor1/2* expression levels between WT and *Gadlor-* KO animals were significant for all organs in which *Gadlor1* and *Gadlor2* were detected. (Heart: *p-value<0.05, Liver and Kidney: ***p-value<0.001, Brain: ****p-value<0.0001). **C.** Experimental design of TAC operation for *Gadlor*-KO and WT littermates. **D.** Heart weight (miligrams) to body weight (grams) ratio (HW/BW) and **E.** lung weight (miligrams) to body weight (grams) ratio of *Gadlor*-KO and WT littermates in sham and after 2 weeks TAC (n≥11). **F.** Representative echocardiography images of *Gadlor*-KO and WT mice before (pre-TAC) and after 2 weeks of TAC (2w TAC) shown in parasternal long axis mode, and analysis of **G.** left ventricle (LV) ejection fraction (EF%), **H.** LV posterior wall thickness in diastole (mm) and **I.** LV end diastolic volume (ul), (n≥12). **J.** Representative Sirius-red fibrosis staining in cardiac tissue sections of sham and 2w TAC animals (scale-bar: 1 mm) and **K.** quantification of fibrotic area (n≥4). **L.** Relative expression of fibrosis marker genes *Col1a1* and *Col3a1* and **M.** hypertrophy marker genes *Nppa, Nppb* and *Mhy7/6* ratio in mRNA levels in heart tissue samples of sham (n=3) and 2w TAC (n=5). **N.** Representative immunofluorescence images of cardiac tissue samples stained with WGA (wheat-germ agglutinin coupled with Alexa Fluor 555) and IB4 (Isolectin B4 coupled with Alexa Fluor 488) in sham and 2-week TAC samples from *Gadlor-*KO and WT littermates, scale-bar: 100 µm. Quantification of **O.** cardiomyocyte cross-sectional area and **P.** myocardial capillary density per cardiomyocyte ratio. Data are shown as mean±SEM. Data normality was evaluated with Shapiro-Wilk test and p-values were calculated with Student’s t-test for comparing two groups and two-way ANOVA for grouped analysis followed with Fisher’s LSD post-hoc test. *p-value<0.05, **p-value<0.01, ***p-value<0.001, ****p-value<0.0001.

The ablation of *Gadlor1/2* was confirmed in knock-out animals by qRT-PCR in different organs **(Figure 2B).** We found *Gadlor1* and *Gadlor2* expression in heart, liver, brain, spleen and kidney, with the highest expression in the brain of WT mice **(Figure 2B)**. To study the effect of *Gadlor1* and *Gadlor2* in cardiac remodeling during pressure-overload, WT and *Gadlor-*KO animals were subjected to TAC or sham surgery and subsequently monitored for 2 weeks **(Figure 2C)**. The carotid flow was measured after 2 days of TAC in the right and left common carotid arteries (RCCA and LCCA, respectively) to evaluate the strength of the aortic constriction. The RCCA/LCCA flow ratio increased to a similar degree in both WT and *Gadlor-*KO animals, indicating comparable pressure overload **(Figure S4F-G).** We found a significant increase in heart weight to body weight (HW/BW) ratio after 2 weeks of TAC in both WT and *Gadlor-*KO compared to sham animals, however, less hypertrophy (i.e. a reduced HW/BW ratio) was observed in *Gadlor-*KO mice **(Figure 2D)**. Additionally, heart-failure-induced pulmonary congestion was only observed in WT animals after TAC, whereas *Gadlor-*KO animals were protected **(Figure 2E)**. In echocardiography, *Gadlor-* KO mice showed less cardiac systolic dysfunction, a markedly reduced left ventricle wall thickness (LVPW) and less chamber dilation (measured as left end-diastolic volume, LVEDV) compared to WT mice after TAC **(Figure 2F-I)**.

### Deletion of *Gadlor1* and *Gadlor2* alleviates cardiomyocyte hypertrophy and fibrosis *in vivo*

Sirius red staining revealed strongly diminished myocardial fibrosis in *Gadlor-*KO mice, which was supported by reduced cardiac *Col1a1* and *Col3a1* mRNA levels after TAC **(Figure 2J-L).** A lower expression of *Nppa, Nppb*, a lower *Myh7/Myh6* RNA ratio, and a reduced cardiomyocyte cross-sectional confirmed blunted cardiac hypertrophy in *Gadlor-*KO animals compared to WT littermates after TAC **(Figure 2M-O).** Additionally, Isolectin B4 (IB4) staining of capillaries revealed a substantial increase of capillary density in the myocardium of *Gadlor-*KO mice compared to WT animals after TAC **(Figure 2N, P).** Together, our data showed that deletion of *Gadlor*1/2 protected against cardiac hypertrophy, systolic dysfunction, capillary rarefaction and myocardial fibrosis during pressure overload.

### Overexpression of *Gadlor1* and *Gadlor2* via EVs triggers cardiac dysfunction and fibrosis

To decipher the functional effect of increased *Gadlor1* and *Gadlor2* levels during pressure overload, we administered control EVs and *Gadlor*-enriched EVs to mouse hearts directly before TAC surgery by intra-ventricular injection (and concomitant cross-clamping of the aorta and pulmonary artery distal of the coronary vessels origin, to allow perfusion of the injected EVs through the coronaries). Control EVs were produced by purifying EC-derived EVs from the supernatant of control adenovirus (Ad.βgal) infected C166 mouse ECs, and *Gadlor1* and *Gadlor2* containing EVs were purified from the supernatant of Ad.*Gadlor1* and Ad.*Gadlor2* infected C166 ECs **(Figure 3A)**. We detected significantly higher levels of myocardial *Gadlor1* and *Gadlor2* after 1 week of EV administration **(Figure 3B)**. *Gadlor*-EV treated mice exerted a marked reduction of cardiac systolic function after 1-week and 2-weeks of TAC, as well as increased wall thickness compared to control EV treated animals (two cohorts were examined, cohort I and II, **Figure 3A-E**). Histological analyses by Sirius red staining showed exaggerated myocardial fibrosis in *Gadlor*-EV treated mice, which was confirmed by qPCR of *Col1a1* and *Col3a1* mRNA expression in the myocardium **(Figure 3F-G)**. Furthermore, increased expression of *Nppa* and *Nppb*, as well as a trend towards larger cardiomyocyte cross-sectional area verified increased hypertrophic remodeling in *Gadlor*-EV treated mice **(Figure 3H-J)**. The myocardial capillary density was not changed between experimental groups **(Figure 3K)**. We concluded that overexpression of *Gadlor1* and *Gadlor2* via administration of endothelial-derived EVs aggravated cardiac dysfunction and hypertrophy, and triggered fibrosis during pressure overload.

**Figure 3:**
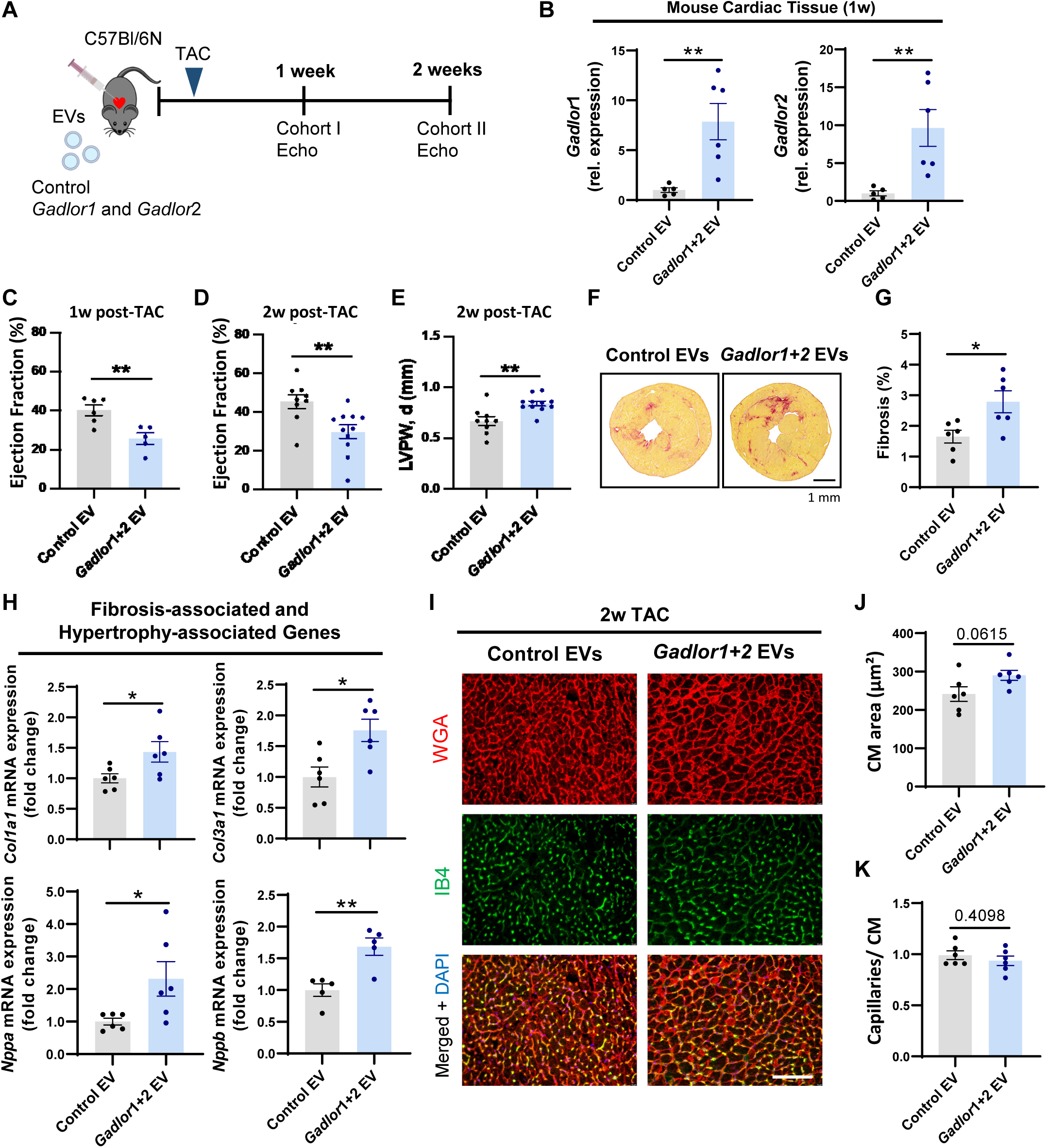
Overexpression of *Gadlor1* and *Gadlor2* via extracellular vesicles triggers cardiac dysfunction and development of fibrosis after TAC. **A.** Scheme of the experimental design depicting the injection of *Gadlor1 and Gadlor2* containing extracellular vesicles for overexpression followed by TAC operation. **B.** Relative expression of *Gadlor1 and Gadlor2* in cardiac tissue samples after 1 week TAC (indicated as cohort I) to validate the overexpression after injection of control (Ad.βgal treated samples, n=5) and *Gadlor-* containing (Ad.*Gadlor1* and Ad.*Gadlor2* treated samples, n=6) EVs. Echocardiographic analysis of **C.** left ventricle (LV) ejection fraction (EF%) after 1 week TAC, and **D.** LV-EF%, **E.** LV posterior wall thickness in diastole (mm) after 2 weeks TAC in mice injected with control EVs and *Gadlor*1 and *Gadlor2* EVs. **F.** Representative images of Sirius-red staining (scale bar: 1 mm) and **G.** quantification of fibrotic area after 2 weeks TAC in mice injected with control EVs and *Gadlor*1 and *2* EVs. **H.** Relative expression of fibrosis marker genes *Col1a1* and *Col3a1* and hypertrophy marker genes *Nppa* and *Nppb* in heart tissue samples of control EV (n=6) and *Gadlor* EVs (n=6) treated mice. **I.** Representative immunofluorescence images of cardiac tissue samples stained with WGA (wheat-germ agglutinin coupled with Alexa Fluor 555) and IB4 (Isolectin B4 coupled with Alexa Fluor 488) in samples from control-EV and *Gadlor*-EV treated animals (scale-bar: 100 µm). Quantification of **J.** cardiomyocyte cross-sectional area and **K.** myocardial capillarization after EV treatment followed by 2 weeks of TAC. Data are shown as mean±SEM. Data normality was evaluated with Shapiro-Wilk test and p-values were calculated with Student’s t-test. *p-value<0.05, **p-value<0.01, ***p-value<0.001, ****p-value<0.0001.

### *Gadlor*-KO mice exert increased mortality during chronic pressure overload

Next, we studied whether the protective effects of *Gadlor1/2* ablation were maintained during persisting long-term pressure overload **(Figure 4A)**. Similar to our findings after 2 weeks of TAC, *Gadlor-*KO animals had a significantly reduced HW/BW ratio after 8-weeks of TAC compared to WT littermates, an improved systolic left ventricular function with reduced posterior wall thickness, an increased capillary density, diminished cardiomyocyte hypertrophy and markedly less fibrosis **(Figure 4B-J)**. Despite retaining better systolic function and improved remodeling features, we observed a considerably higher mortality rate in *Gadlor*-KO mice starting between two and three weeks after TAC **(Figure 4K)**. Interestingly, WT and KO animals were indistinguishable in terms of appearance, behavior and mobility after TAC operation, and death in *Gadlor-*KO mice was unexpected and sudden in nature, possibly due to cardiac arrhythmia.

**Figure 4:**
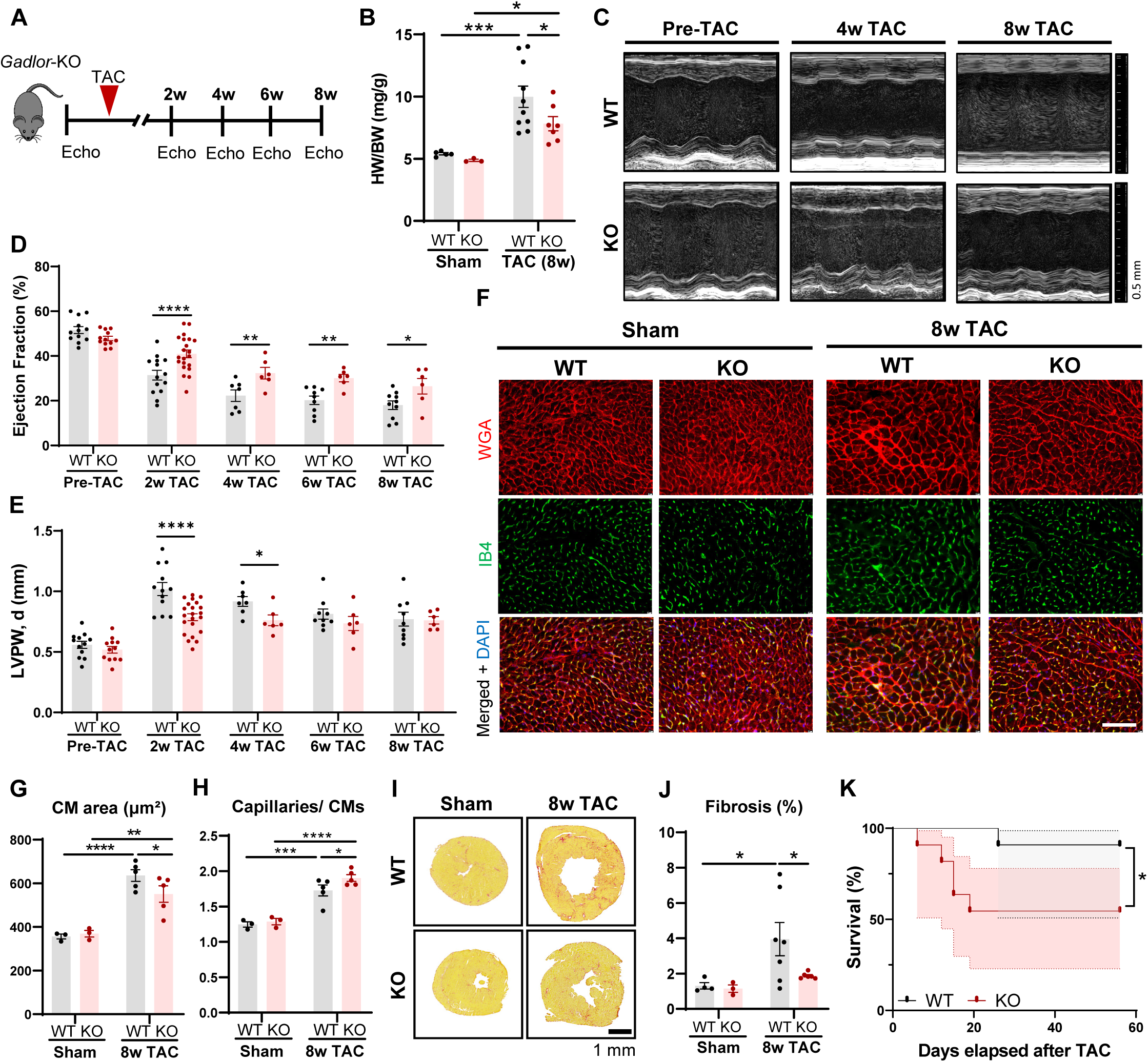
*Gadlor*-KO mice showed higher mortality despite retaining better structural and physiological remodeling after persisting pressure-overload. **A.** Experimental design of long-term TAC operation for *Gadlor*-KO and WT littermates up to 8-weeks after TAC. **B.** Heart weight (mg) to body weight (g) ratio (HW/BW) of *Gadlor*-KO and WT mice in age comparable sham (n≥3) and after 8-weeks TAC (n≥7). **C.** Representative echocardiography images of *Gadlor*-KO and WT mice before (pre-TAC) and after 4-weeks and 8-weels TAC shown in parasternal long axis mode, and analysis of **D.** left ventricle (LV) ejection fraction (EF%) and **E.** left ventricle (LV) posterior wall thickness in diastole (mm), (unpaired analysis)**. F.** Representative immunofluorescence images of cardiac tissue samples stained with WGA (wheat-germ agglutinin coupled with Alexa Fluor 555) and IB4 (Isolectin B4 coupled with Alexa Fluor 488) in sham (n=3) and 8-week TAC (n=5) samples from *Gadlor-*KO and WT littermates, scale-bar: 100 µm. Quantification of **G.** cardiomyocyte cross-sectional area and **H.** myocardial capillary density per cardiomyocyte ratio. **I.** Representative Sirius-red fibrosis staining in cardiac tissue sections of sham (n≥3) and 8-week TAC (n≥6) mice (scale-bar: 1 mm) and **J.** quantification of fibrotic area (%). **K.** Survival of *Gadlor-*KO and WT mice after TAC (n=11). Probability of survival compared with log-rank (Mantel-Cox) test. Data are shown as mean±SEM. Data normality was evaluated with Shapiro-Wilk test and p-values were calculated with Student’s t-test for comparing two groups and two-way ANOVA for grouped analysis followed with Fisher’s LSD post-hoc test. *p-value<0.05, **p-value<0.01, ***p-value<0.001, ****p-value<0.0001.

### *Gadlor1* and *Gadlor2* affect gene expression and angiogenic function in ECs

Since ECs are the main cardiac cell-type expressing *Gadlor1/2*, and since we observed an increased capillary density in the myocardium of *Gadlor*-KO mice after two and 8 weeks of TAC (see above), we performed RNA sequencing from isolated WT and *Gadlor-*KO ECs after two weeks of TAC. The purity of ECs in our MACS-based isolation procedure is >90%.^34^ The genome-wide transcriptome analysis via bulk RNAseq identified a total of 3355 differentially expressed (DE) genes between WT and *Gadlor-*KO, which are shown in the heatmap **(Figure 5A)**. Gene ontology analysis from DE genes and RT-qPCR revealed and confirmed upregulation of angiogenesis, mitotic cell cycle and respiratory electron transport and downregulation of inflammatory response genes in *Gadlor*-KO EC after two weeks of TAC **(Figure 5B-D)**. In addition, myocardial staining for Ki67 as a proliferation marker revealed more Ki67-positive EC in *Gadlor-* KO mice after TAC **(Figure 5E-F)**. In turn, overexpression of *Gadlor1* and *Gadlor2* by adenovirus in C166 mouse ECs blunted the increase in EC sprouting activity during FGF2 stimulation and reduced sprouting further in the presence of TGFβ in comparison to Ad.β*gal* (control) infected cells **(Figure 5G-H)**. In summary, *Gadlor* lncRNAs exert anti-angiogenic properties in ECs.

**Figure 5:**
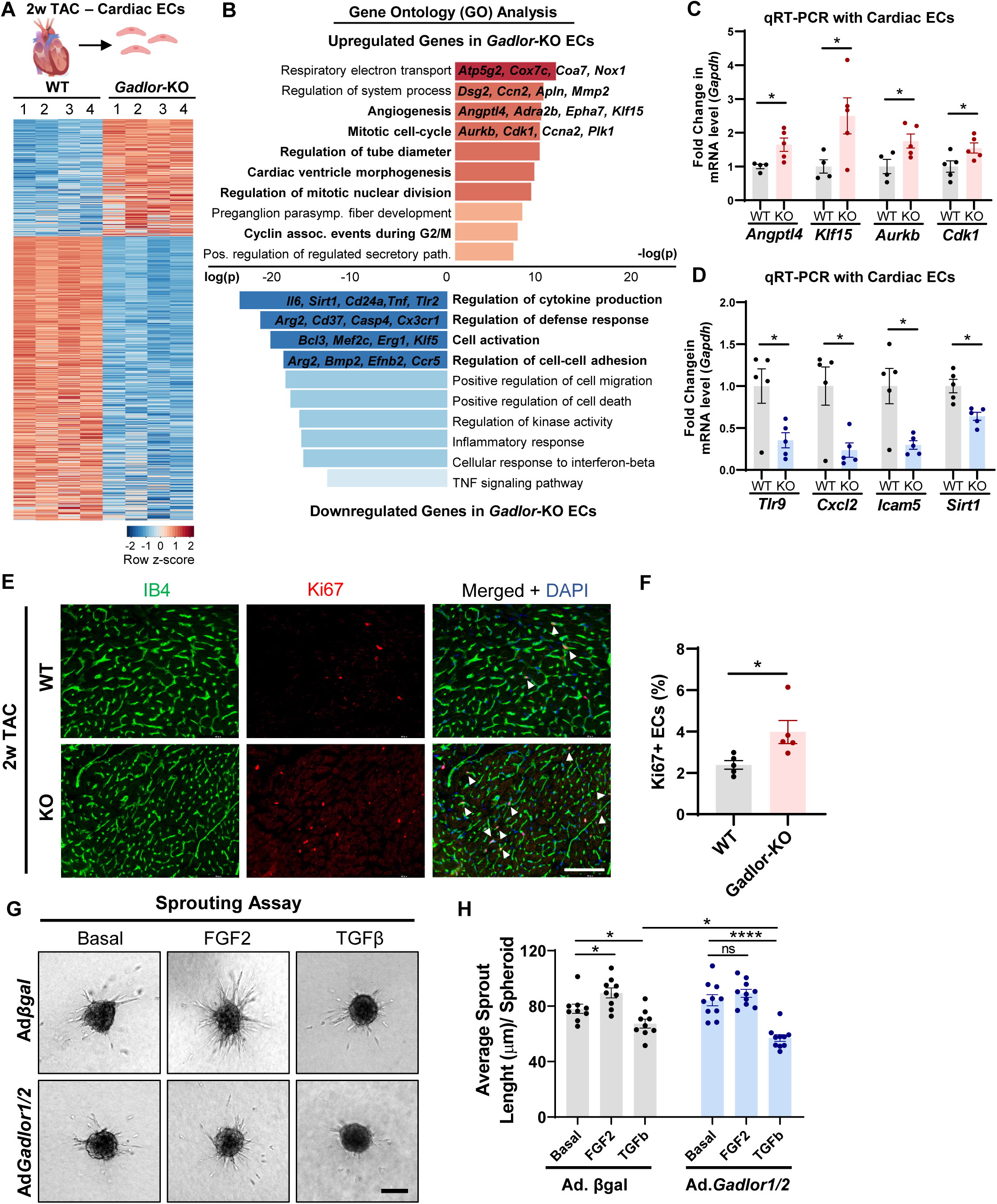
Deletion of *Gadlor* lncRNAs induces angiogenesis and expression of cell cycle genes in cardiac ECs during pressure-overload. **A.** Heatmap showing differentially regulated genes revealed by bulk RNAseq of cardiac endothelial-cells after 2-weeks TAC. **B.** Bar plots showing the gene-ontology (GO) analysis of upregulated and downregulated genes in *Gadlor-*KO ECs compared to WT ECs after 2-weeks TAC (Red: Upregulated in *Gadlor-*KO, Blue: Downregulated in *Gadlor-*KO). Some exemplary genes were listed for selected GO-terms. **C-D.** Validation of selected genes from RNAseq data with qRT-PCR in isolated cardiac ECs after 2 weeks of TAC (Red: Upregulated in *Gadlor-*KO, Blue: Downregulated in *Gadlor-*KO). **E.** Representative immunofluorescence images of cardiac tissue samples stained with IB4 (Isolectin B4 coupled with Alexa Fluor 488) and Ki67 (coupled with Alexa Fluor 555) in 2-week TAC samples from *Gadlor-*KO and WT littermates, Scale bar: 100 µm. (Arrows showing Ki67+ ECs) **F.** Quantification of Ki67 positive endothelial-cells percentage (Ki67+ ECs%). **G.** Representative images of sprouting assay with C166 mouse ECs after adenovirus treatment (Ad*βgal* and Ad*Gadlor1/2*) followed by 24-hour FGF2 or TGFβ treatment. 3-dimentional (3D) collagen matrix embedded spheroids were allowed to form sprouts for 24 hours. Scale bar: 100 μm. **H.** Quantification of average sprout length per spheroid. Data are shown as mean±SEM. Data normality was evaluated with Shapiro-Wilk test. P-values were calculated with Student’s t-test for comparing two groups. For grouped analysis, p-values were evaluated with two-way ANOVA followed with Fisher’s LSD post-doc test. *p-value<0.05, **p-value<0.01, ***p-value<0.001, ****p-value<0.0001.

### Cardiac FB of *Gadlor*-KO mice induce less fibrosis-associated genes after TAC

Based on our findings, cardiac fibroblasts (FBs) exerted the second highest *Gadlor* lncRNAs expression after ECs in the myocardium **(Figure 1C).** To study the reason for decreased fibrosis in hearts upon *Gadlor* knock-out after TAC, we performed bulk RNAseq from isolated cardiac FBs after two weeks of TAC. The purity of cardiac FBs in our MACS-based isolation procedure is >90%.^34^ Genome-wide analysis revealed a total of 915 downregulated and 998 upregulated genes in *Gadlor-*KO FBs compared to WT mice after TAC, which are shown in a heatmap in **Figure S5A**. Amongst these, we detected a strong downregulation of mainly ECM organization and growth factor associated genes (confirmed by RT-qPCR: *Fgfr1, Igf1, Col1a1, Col3a1, Col6a1*) **(Figure S5B-D)**. Additionally, genes related to the regulation of angiogenesis, blood vessel and endothelium development were upregulated in *Gadlor-*KO FBs, which suggested a contribution of FBs to the enhanced angiogenesis observed in *Gadlor-*KO mice upon overload. The increased expression of *Angpt2, Efna1, Aurkb, Dll1* in cardiac FBs from *Gadlor KO* mice was confirmed by RT-qPCR **(Figure S5C)**. Upon adenoviral overexpression of *Gadlor1* and *Gadlor2* followed with phenylephrine (PE) stimulation, neonatal rat cardiac fibroblasts (NRFB), in turn, exerted a decreased expression of *Angpt2*, and an increased expression of *Col1a1* and *Col3a1*, but no differential expression was observed for *Col4a1* or *Col6a1* **(Figure S5E)**. We concluded that *Gadlor* lncRNAs trigger pro-fibrotic gene expression in cardiac FBs.

### *Gadlor* lncRNAs are transferred to cardiomyocytes via EC-derived EVs

Since *Gadlor1* and *Gadlor2* were mainly secreted by ECs, we reasoned that these lncRNAs might exert paracrine effects due to EV-mediated transfer into cardiomyocytes, where they could affect contraction, arrhythmia, or hypertrophy. To analyze whether *Gadlor*1/2 are transferred from ECs to cardiomyocytes, we initially overexpressed *Gadlor1*/*2* by adenovirus in C166 mouse ECs, and purified EVs from the supernatant of these cells **(Figure 6A)**. We confirmed the enhancement of *Gadlor1/2* expression in EVs upon *Gadlor1/2* overexpression versus control EVs **(Figure 6B)**. The collected EVs were transferred either to neonatal rat cardiomyocytes (NRCMs) or fibroblasts (NRFBs). RT-qPCR analysis showed a significantly higher abundance of *Gadlor1/2* in NRCMs incubated with *Gadlor*-EVs compared to samples incubated with control-EVs or cardiomyocytes without any EV treatment, which indicated that EVs were taken up by cardiomyocytes **(Figure 6C)**. *Gadlor* lncRNAs were also transferred from EC to cardiomyocytes by control EVs, since higher cardiomyocyte *Gadlor1/2* levels were observed after control EV compared to non-EV treatment. Additionally, a mild increase in both *Gadlor1* and *Gadlor2* abundance was also observed in NRFBs after incubation with *Gadlor* EVs, however, at a much lower degree compared to NRCMs **(Figure 6C)**. We visualized the uptake of the fluorescently labelled (PKH67) EVs into recipient cardiomyocytes, which was analyzed with confocal microscopy after 6 hours and 24 hours **(Figure 6D)**. 3D imaging by confocal Z-stacks confirmed that EVs were internalized into the recipient cardiomyocytes rather than being attached to the surface **(Supplemental Video 1)**.

**Figure 6:**
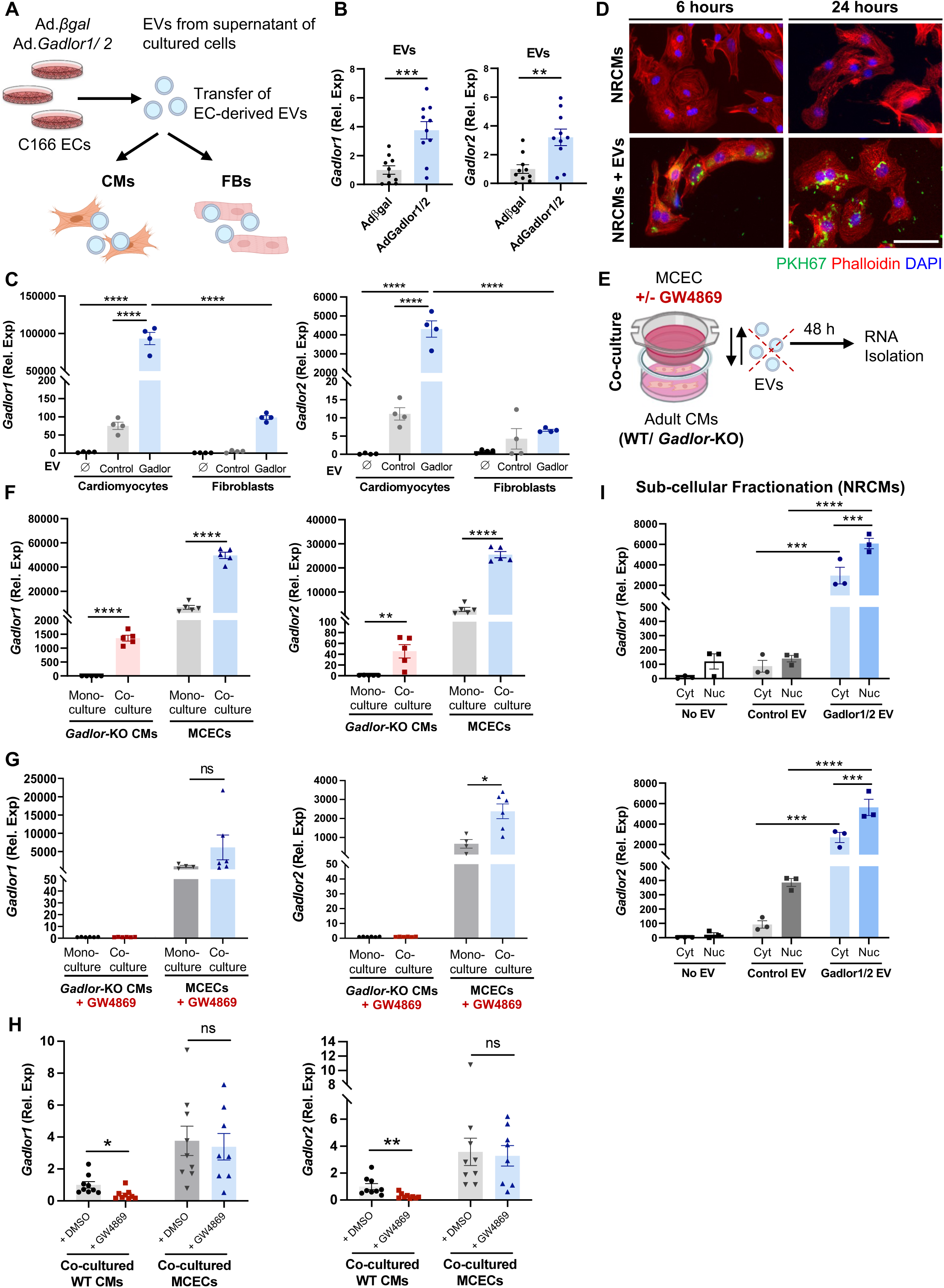
Secreted *Gadlor* lncRNAs are transferred to cardiomyocytes via endothelial-derived EVs. **A.** Schematic representation of experimental design of EV (isolated from C166 endothelial cells, EC, supernatant) transfer to NRCMs or NRFBs. **B.** Validation of *Gadlor1* and *Gadlor2* overexpression in isolated EVs. **C.** RT-qPCR detection of *Gadlor1* and *Gadlor2* after transferring control EVs (isolated from Adβgal treated ECs) or Gadlor-EVs (isolated from Ad*Gadlor1/2* treated ECs) on neonatal rat cardiomyocytes (NRCMs) or neonatal rat cardiac fibroblasts (NRFBs). The samples were collected after 6 hours of EV incubation. **D.** Visualization of NRCMs that were incubated with PKH67-labelled (green) EVs isolated from C166 ECs for 6 hours and 24 hours. Staining was performed with Phalloidin (red) and DAPI (blue). Scale bar: 50 μm. **E.** Scheme of co-culture experiment with transwell (1 μm pore diameter – allows EV transfer). Mouse WT cardiac EC (MCEC) on top cultured with *Gadlor*-KO or WT CMs in the bottom well without direct cell contact. **F.** RT-qPCR detection of *Gadlor1* and *Gadlor2* in isolated *Gadlor-*KO CMs and MCEC mouse ECs after 48 hours of mono-culture or co-culture as indicated. **G-H.** RT-qPCR detection of *Gadlor1* and *Gadlor2* in isolated adult *Gadlor-*KO CMs (G) or WT CMs (H) and MCECs that were incubated with GW4869 as indicated. Samples were collected after 48 hours of mono-culture and co-culture. DMSO was used as vehicle control. **I.** *Gadlor1* and *Gadlor2* expression in the cytosol and nucleus of the recipient NRCMs after EV transfer (as shown in A). Data are shown as mean±SEM. Data normality was evaluated with Shapiro-Wilk test and p-values were calculated with Student’s t-test or one-way ANOVA with Fisher’s LSD post-hoc test were applied for comparing two or multiple groups, respectively. Two-way ANOVA with Fisher’s LSD post-hoc test was applied for grouped analysis when applicable. *p-value<0.05, **p-value<0.01, ***p-value<0.001, ****p-value<0.0001.

Furthermore, we performed co-culture experiments with isolated *Gadlor*-KO cardiomyocytes (lacking endogenous *Gadlor1/2* expression) and WT mouse cardiac endothelial cells (MCEC, on cell culture inserts, which allow the transfer of EVs, due to a pore size of 1 μm) for 48 hours **(Figure 6E)**. *Gadlor*-KO cardiomyocytes were used to exclude confounding effects of endogenous *Gadlor1/2* expression with exogenous *Gadlor1/2* uptake from EC-derived EVs. As expected, we did not find any *Gadlor1/2* expression in isolated *Gadlor*-KO cardiomyocytes when they were kept in monoculture, whereas we detected *Gadlor1* and *Gadlor2* after co-culture with WT MCECs **(Figure 6F)**. This suggested that *Gadlor1/2* were transferred into *Gadlor-*KO cardiomyocytes from WT MCECs, possibly via EV-mediated cell-to-cell transfer. MCECs that were co-cultured with *Gadlor*-KO cardiomyocytes also exerted an increase of *Gadlor1/2* expression **(Figure 6F)**, which might suggest a possible feedback mechanism between cardiomyocytes and ECs. The addition of GW4869 as inhibitor of exosome (i.e. small EV) generation into the co-culture system ablated the transfer of EC derived *Gadlor1/2* to KO cardiomyocytes as well as the putative feedback mechanisms, suggesting that both might depend on small EVs **(Figure 6G)**. Next, we wanted to assess whether WT cardiomyocyte depend on ECs for *Gadlor1/2* expression. Indeed, inclusion of GW4869 into co-culture of WT cardiomyocytes with WT MCECs, markedly reduced *Gadlor1/2* levels in the cardiomyocytes. This indicated that ECs provide *Gadlor1/2* to cardiomyocytes and might thereby relay paracrine signals to these cells **(Figure 6H)**.

Sub-cellular fractionation studies to localize *Gadlor* lncRNAs following EV-mediated transfer showed that *Gadlor1* and *Gadlor2* were detected in the cytosol and in a bit higher level in the nucleus of recipient cardiomyocytes **(Figure 6I)**. Overall, endothelial-derived *Gadlor1/2* containing EVs are mainly taken up by cardiomyocytes, where they might act in the cytosol as well as in the nucleus. Cardiac FBs may mainly rely on their own cellular *Gadlor1/2* expression.

### *Gadlor* lncRNAs bind to calcium/calmodulin-dependent protein kinase type II in cardiomyocytes

LncRNAs can interact with proteins to affect their regulatory functions.^35^ Even though cardiomyocytes showed the least endogenous levels of *Gadlor* lncRNAs, we still observed strong effects in cardiomyocytes (e.g. reduced hypertrophy and potentially arrhythmia) after *Gadlor1/2* deletion upon pressure overload. To decipher this further, we aimed at identifying the binding partners of *Gadlor1/2* in cardiomyocytes by performing RNA antisense purification coupled with mass-spectrometry (RAP-MS) in HL1 cardiac muscle cells after overexpression of *Gadlor1/2* **(Figure 7A)**. Interestingly, we found calcium/calmodulin-dependent protein kinase type II, delta (CaMKIIδ) as the most significant binding partner of *Gadlor1/2* **(Figure 7B)**. Next, we confirmed the interaction between CaMKII and *Gadlor1/2* with native RNA immunoprecipitation (RIP), i.e. immunoprecipitation of CaMKII, followed by qRT-PCR to detect *Gadlor1* and *Gadlor2* **(Figure 7C)**. We then investigated whether *Gadlor1/2* modulates CaMKII function by assessing phospholamban Threonine-17 (Thr17) phosphorylation, which is mediated directly by CaMKII. Indeed, we found reduced and increased Thr17 phospholamban phosphorylation in *Gadlor-*KO mice or *Gadlor*-EV treated mice after TAC, respectively **(Figure 7D-G)**. This indicated that *Gadlor1/2* might promote CaMKII activation in cardiomyocytes.

**Figure 7.**
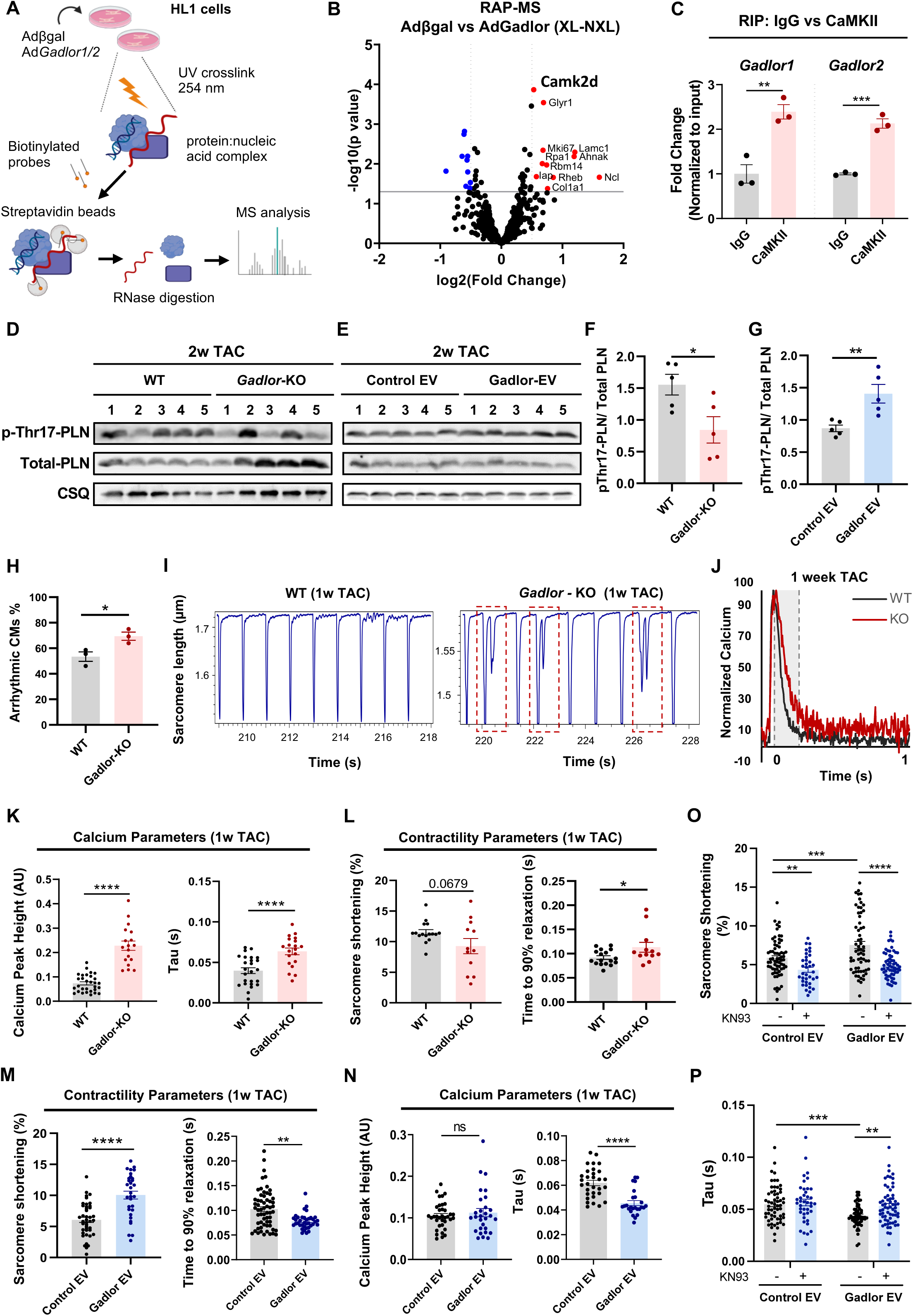
*Gadlor* lncRNAs bind to calcium/calmodulin-dependent protein kinase type II in cardiomyocytes. **A.** Experimental scheme of RNA-antisense purification coupled with mass spectroscopy (RAP-MS) to identify specific interaction partners of Gadlor1/2 in HL1 cardiomyocytes treated with βgal or Gadlor1/2 adenovirus. **B.** Volcano plot indicating the identified interaction partners of Gadlor1 and Gadlor2 in cardiomyocytes with RAP-MS (Fold change enrichment (FC) threshold 1.2 and p-value < 0.05). UV crosslinking was used to stabilize protein:RNA interactions and non-crosslinked (N-XL) samples were used as background and removed from crosslinked (XL) samples for each indicated condition. **C.** RNA immunoprecipitation (RIP) performed with anti-IgG or anti-CaMKII antibodies from HL-1 cardiac muscle cell lysate to validate the interaction with Gadlors. RT-qPCR analysis showing the enrichment of Gadlor1 and Gadlor2 compared to IgG controls. Data was normalized to input RNA expression. (n=3 for each condition) **D-E.** Exemplary Western blots of phospho-Thr17-PLN and total PLN with CSQ as loading control in (D) WT and Gadlor-KO and (E) in control-EV and Gadlor-EV treated mouse heart tissue samples after 2 weeks TAC**. F-G.** Quantification of Western-blots. **H.** Percentage of isolated adult WT and Gadlor-KO cardiomyocytes with arrhythmic phenotype after 1-week TAC recorded by Multicell HT system (Ion-Optix) Cell number≥30 in each condition. **I.** Exemplary traces of sarcomere shortening (contractility) of adult WT and *Gadlor-*KO cardiomyocytes isolated after 1 week of TAC (1w TAC). Images showing the measurement of transients in WT-CMs in which no arrhythmia was detected, and detection of arrhythmic beating showed as a secondary peak indicating an after-contraction before the end of the first contraction has been reached in GADLOR-KO-CMs. Recordings of transients (nine transient in each measurement cycle) are obtained from a single cardiomyocyte. Images were analysed with the CytoSolver Transient Analysis Tool. **J – N.** Sarcomere contractility and calcium transient measurements were recorded in parallel with Multicell HT system with continuous perfusion of basal media on isolated adult CMs after 1-week TAC. **J.** Representative calcium transient recordings of WT and Gadlor-KO mice shown as normalized levels of calcium (%). **K.** Calcium parameters including calcium peak height (arbitrary units, AU) and decay time (tau, seconds). **L.** Contractility parameters including sarcomere shortening (%) and relaxation. **M.** Sarcomere function and **N.** calcium transient and decay time constant (tau). Recordings were obtained from adult cardiomyocytes isolated from WT mice after 1 week TAC followed by incubation of control-EV (Ad.β*gal*) and *Gadlor1/2*-EVs (Ad.*Gadlor1* and Ad.*Gadlor2*) for 4-6 hours. **O-P.** Sarcomere contractility and calcium tau measurements in cardiomyocytes as described before with and without addition of the CamKII inhibitor KN93. Data are shown as mean±SEM. Data normality was evaluated with Shapiro-Wilk test and p-values were calculated with Student’s t-test (Mann-Whitney for non-parametric) for comparing two groups. *p-value<0.05, **p-value<0.01, ***p-value<0.001, ****pvalue<0.0001.

Since *Gadlor* lncRNAs had influenced cardiac function, and CaMKII was identified as binding partner, which has a known role in cardiomyocyte excitation-contraction coupling, we investigated cardiomyocyte contractility and calcium dynamics with the CytoCypher High-Throughput (HT) system (Ionoptix) in isolated adult cardiomyocytes after one week of TAC. Interestingly, during measurements, a considerably higher number of *Gadlor*-KO cardiomyocytes with arrhythmic behavior was detected compared to WT cardiomyocytes **(Figure 7H)**. Sarcomeric length traces **(Figure 7I)** indicated that *Gadlor*-KO cardiomyocytes exhibited double contractions even though they were paced with electrical field stimulation, in which each stimulation usually results in a single contraction.

Interestingly, *Gadlor-*KO cardiomyocytes displayed a much slower re-uptake of calcium into the sarcoplasmic reticulum (SR), which led to prolonged calcium transients. We also observed an increased calcium peak height in *Gadlor-*KO cardiomyocytes **(Figure 7J-K).** Additionally, while sarcomere shortening and its timing were not significantly affected, cardiomyocyte relaxation was delayed **(Figure 7L)**. Treatment of adult mouse cardiomyocytes with *Gadlor1/2* enriched versus control EC-derived EVs, in turn, led to augmented sarcomere shortening and especially faster cardiomyocyte relaxation **(Figure 7M)**. While calcium peak height was not changed, a faster diastolic calcium re-uptake was detected by a decreased tau value **(Figure 7N)**. Collectively, *Gadlor-*KO myocytes displayed an enhanced propensity to arrhythmic beating, slower diastolic relaxation and slower calcium re-uptake into SR, which entailed increased intracellular calcium levels. Overexpression of *Gadlors1/2,* in turn, led to increased sarcomere shortening, faster cardiomyocyte relaxation and faster re-uptake of calcium into the SR. These effects were partially inhibited upon KN93 (CaMKII inhibitor) treatment **(Figure 7O)**, which decreased the sarcomere shortening and increased the time needed for re-uptake of calcium into the SR (tau) in *Gadlor-*EV treated cardiomyocytes, indicating that *Gadlor1/2* exert their effects on excitation-contraction coupling at least in part through CaMKII activation.

### *Gadlor1* and *Gadlor2* promote pro-hypertrophic and mitochondrial gene-expression in cardiomyocytes

Since *Gadlor1/2* were in part localized in cardiomyocyte nuclei after their transfer **(Figure 6I)**, we hypothesized that they might also affect gene expression in these cells. Comprehensive transcriptome analysis after two weeks of TAC between isolated WT and *Gadlor-*KO cardiomyocytes revealed that 2492 genes were up-, while 2956 were downregulated after ablation of *Gadlor1/2* as shown in the heatmap in **Figure 8A**. Functional annotation of DE genes showed that genes related to vasculature development, ECM organization and regulation of cytokine production were upregulated **(Figure 8B)**, while mitochondrial organization, TCA cycle and cardiac muscle contraction/hypertrophy related genes were downregulated in *Gadlor*-KO cardiomyocytes **(Figure 8B)**. We confirmed the upregulation of *Il6, Tlr9, Col15a1, Col14a1, Adam8* mRNAs, and the downregulation of *Camk2d, Gata4, Actn2, Mfn2* mRNAs in Gadlor-KO cardiomyocytes after TAC by RT-qPCR **(Figure 8C-D)**. To rule out a reduced number of mitochondria as a reason for the downregulation of mitochondrial genes, we assessed the mitochondrial-to-nuclear DNA ratio, but detected no changes in *Gadlor*-KO versus WT mice, indicating a similar number of mitochondria under both conditions **(Figure 8E)**. We next assessed the effects of *Gadlor1/2* overexpression on cardiomyocyte gene-expression after PE stimulation **(Figure 8F)**. *Gadlor1/2* overexpression in large parts revealed opposite effects of what was found in *Gadlor*-KO cardiomyocytes, for example it promoted the expression of genes assigned to the GO-class “cardiac muscle contraction” (*Gata4*, *Camk2d*, *Rcan1.4*), which mainly exert pro-hypertrophic function in cardiomyocytes. These effects were partially reversed upon treatment with KN93, indicating CaMKII-dependent effects of *Gadlor1/2* on pro-hypertrophic (and other) gene expression.

**Figure 8.**
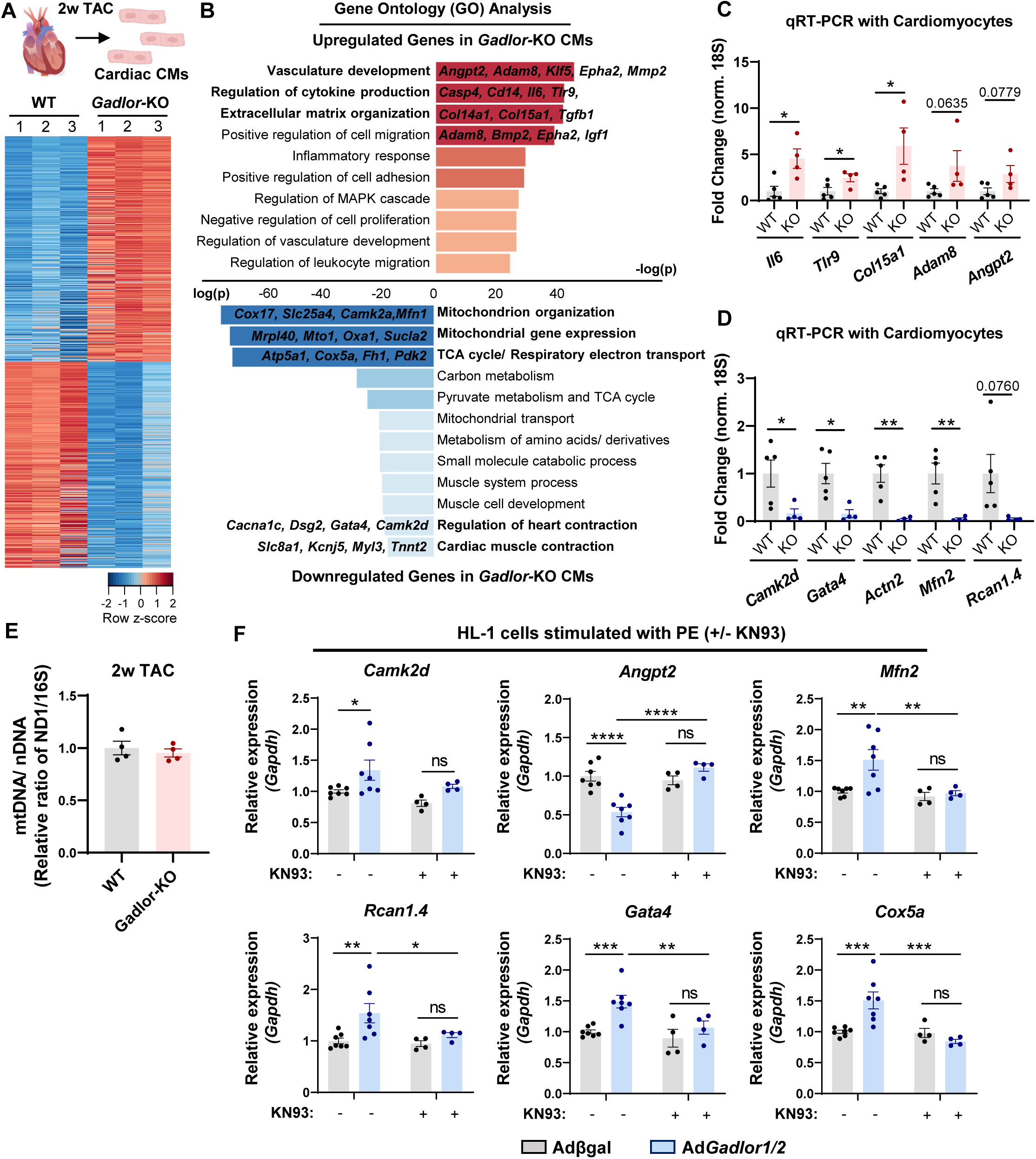
Deletion of *Gadlor1* and *Gadlor2* affects gene expression in cardiomyocytes. **A.** Heatmap showing differentially regulated genes analysed by bulk RNAseq of isolated cardiomyocytes after 2-weeks TAC. **B.** Bar plots showing the gene-ontology (GO) analysis of upregulated and downregulated genes in Gadlor-KO CMs after 2-weeks TAC. (Red: Upregulated in Gadlor-KO, Blue: Downregulated in Gadlor-KO). Exemplary genes were listed for selected GO-terms**. C-D.** Validation of selected upregulated and downregulated genes from RNAseq data with qRT-PCR. **E.** Analysis of mitochondrial DNA (mtDNA) to nuclear DNA (nDNA) ratio determined via quantitative PCR by using WT and Gadlor-KO heart tissue samples isolated 2-weeks after TAC. Assessment of ND1 and 16S DNA abundance was used to determine the mtDNA/ nDNA ratio. **F.** qRT-PCR of selected genes in HL-1 cardiac muscle cells overexpressing Gadlor lncRNAs after Gadlor1/2 adenovirus treatment (βgal as control) followed by phenylephrine (PE) stimulation (100 μM, 24 hours) with and without KN93 (1 μM, 2 hours) addition. Data are shown as mean±SEM. Data normality was evaluated with Shapiro-Wilk test. *p*-values were calculated with Student’s t-test for parametric (or Mann-Whitney for non-parametric) for comparing two groups. Two-way ANOVA with Fisher’s LSD post-hoc test was applied for grouped analysis. ns: no significance, *p-value<0.05, **p-value<0.01, ***p-value<0.001, ****p-value<0.0001.

Comparing the effects of *Gadlor* knock-out in the different cardiac cell types after TAC revealed a strong overlap among upregulated and downregulated genes in FBs, ECs and cardiomyocytes **(Figure S6)**. For example, *Gadlor* knock-out led to the upregulation of energy-coupled proton transport mitochondrial genes, actin filament associated genes, and mitotic cell cycle related genes in all three cell types, in addition to the upregulation of cardiac muscle contraction and blood vessel morphogenesis genes in two cell types. On the other hand, genes related to the negative regulation of cell differentiation, cartilage development and the regulation of membrane potential were downregulated in all three cell types, while genes related to cell migration, inflammation, extracellular matrix organization, heart contraction, calcineurin/NFAT signaling and mitochondrial organization were downregulated in at least two of the cell types **(Figure S6)**. Therefore, *Gadlor1/2* effects on gene expression contain common, but also unique targets in different cardiac cell types. The findings of this study are graphically summarized in **Figure S7**.

## Discussion

Here, we describe two lncRNAs (termed *Gadlor1* and *Gadlor2*) as central regulators of cardiac failure during pressure overload. *Gadlor1* and *Gadlor2* were jointly upregulated in mouse and human failing hearts. In the serum of patients with aortic stenosis, high *GADLOR2* levels were associated with a low ejection fraction. Accordingly, enhanced experimental *Gadlor1/2* expression in mouse hearts triggered aggravated heart failure, myocardial hypertrophy and fibrosis, while combined *Gadlor*-KO mice were protected from cardiac dysfunction, fibrosis and vascular rarefaction during short and persisting pressure overload. Paradoxically, despite retaining better heart function and remodeling features, the complete lack of *Gadlor* lncRNAs during chronic, persisting pressure overload entailed sudden death, most likely due to arrhythmia.

Among the different cell types in the myocardium, we found the highest *Gadlor1/2* expression in ECs, where reduced *Gadlor1/2* levels entailed an increased capillary density and cell proliferation after TAC. Importantly, enhanced myocardial angiogenesis was reported to promote cardiac function during pressure overload.^20, 22^ *Gadlor1/2* overexpression in isolated ECs reduced angiogenic activity, indicating that *Gadlor* lncRNAs inhibit angiogenesis in ECs in a cell autonomous manner. Because lncRNAs often exert their effects by modifying gene expression, we performed transcriptome profiling by RNA sequencing in cardiac ECs after TAC. Indeed, we found that upon *Gadlor1/2* deletion, EC upregulate genes related to angiogenesis, cell cycle and mitosis. Besides this cell-autonomous role, *Gadlor*-KO mice also exerted an increased expression of proangiogenic genes in cardiomyocytes and FBs during TAC, which are both known to affect angiogenesis in a paracrine manner.^20, 21, 36, 37^

In addition to ECs, cardiac FBs exhibited high levels of *Gadlor1/2* expression in the heart. Transcriptomic profiling in cardiac FBs revealed the downregulation of many extracellular matrix-organizing genes, including pro-fibrotic growth factors, their receptors as well as matrix proteins themselves in *Gadlor*-KO mice. Our data imply that the pro-fibrotic role of *Gadlor1* and *Gadlor2* are mediated by their direct effect on gene expression in cardiac FBs. Increased and decreased fibrosis will likely play a major role in reduced and improved heart function during TAC in mice with enhanced or ablated *Gadlor* expression, respectively.

Remarkably, *Gadlor1/2* are mainly secreted within EVs from ECs. This finding indicated a potential paracrine role of *Gadlor* lncRNAs in the heart. Indeed, EVs were reported to contain mainly miRNAs, but also lncRNAs and mRNAs, and to protect these RNA species from degradation by extracellular RNases.^38^ EVs might thereby play a role in intercellular communication, especially during stress stimulation or disease, when their production is typically upregulated.^39^ In cell culture experiments, we found an effective uptake of *Gadlor1/2* from endothelial-derived EVs in neonatal and adult cardiomyocytes. When adult cardiomyocytes from *Gadlor*-KO mice that naturally could not upregulate intrinsic *Gadlor1/2* lncRNA expression were co-cultured in a two-chamber system (allowing the transfer of EVs, but not cells between the compartments) with wild-type ECs, these ECs in the upper well increased *Gadlor1/2* levels in the knock-out cardiomyocytes at the bottom well, an effect that was blunted by inhibition of small EV generation. In addition, the co-cultured WT ECs markedly upregulated their *Gadlor1/2* expression compared to when they were cultured alone, which was also blunted by EV inhibition. Perhaps even more importantly, when WT ECs and WT cardiomyocytes were co-cultured, EV-inhibition strongly reduced *Gadlor1/2* levels in the WT cardiomyocytes. This demonstrated that EC-derived EVs are crucial to provide *Gadlor1/2* to cardiomyocytes, in which they regulate calcium cycling as well as gene-expression. Our results therefore indicated a reciprocal intercellular signaling circuit between ECs and cardiomyocytes, whereby ECs provide *Gadlor* lncRNAs within EVs to cardiomyocytes, which signal back to ECs by currently unknown (perhaps EV based) mechanisms to indicate insufficient *Gadlor1/2* levels, when necessary. Indeed, signal responsive EV release had been previously described.^39^ In conclusion, *Gadlor1/2* exert a strong effect in intercellular communication in the heart during pathological overload, on one hand by directly serving as intercellular messenger between ECs and cardiomyocytes, and on the other hand by regulating the gene-expression of secreted growth factors in cardiomyocytes, ECs and FBs, thereby affecting, for instance, angiogenesis and fibrosis.

In cardiomyocytes, *Gadlor1/2* lncRNAs are taken up from ECs and localize to the cytosolic as well as to the nuclear compartment. Accordingly, we identified CaMKII as binding partner of *Gadlor1/2* in cardiomyocytes, which localizes to both compartments and is activated mainly by calcium/calmodulin, autophosphorylation and oxidation or S-nitrosylation.^40^ Its activity is known to be about 3-fold upregulated in failing human hearts.^41^ In cardiomyocytes, CaMKII promotes on one hand maladaptive features such as hypertrophy and fibrosis by phosphorylating HDAC4, leading to de-repression of MEF2-dependent, pro-hypertrophic gene expression.^42, 43^ On the other hand, it has adaptive functions, for example, it promotes calcium re-uptake into the SR by phosphorylating phospholamban at Thr17.^44–46^ Because *Gadlor1/2* bind CaMKII and *Gadlor*-KO mice exerted reduced, while *Gadlor*1/2 overexpression led to enhanced Thr17 phosphorylation, we concluded that the *Gadlor* lncRNAs might promote CaMKII activation in a feed-forward loop in cardiomyocytes during pathological overload. Decreased CaMKII activity could account for the improved cardiac remodeling features (i.e. reduced hypertrophy and improved systolic heart function) in *Gadlor*-KO mice, but at the same time slow down calcium re-uptake into the SR, leading to increased cytosolic calcium levels with pro-arrhythmic activity. Indeed, upon *Gadlor1/2* overexpression, pharmacological CaMKII inhibition partially reversed pro-hypertrophic gene expression and reversed increased calcium cycling in isolated cardiomyocytes. Besides facilitating CaMKII activity by direct protein binding, *Gadlor1/2* also appear to promote *Camk2d* expression at the mRNA level. As another pro-hypertrophic gene, *Gata4* is also positively regulated by *Gadlor1/2* in cardiomyocytes. Although details how *Gadlor1/2* directly regulate gene-expression in the cardiomyocyte nucleus will have to be addressed in future studies, this could work through binding to the chromatin reader protein GLYR1 (also known as NDF, NPAC or NP60), which we also found as direct binding partner of *Gadlor1/2* and that is strongly involved in the regulation of cardiomyocyte gene-expression during differentiation by binding to the gene body of actively transcribed genes, often together with GATA4.^47^

In conclusion, upregulation of Gadlor1/2 in failing hearts and their transfer into cardiomyocytes might be a compensatory mechanism aiming at promoting calcium re-uptake into the sarcoplasmic reticulum during hemodynamic overload, although this comes at the cost of aggravated hypertrophy and fibrosis. Considering their dichotomous role, *Gadlor1/2* inhibition in cardiomyocytes might therefore not be a suitable therapeutic strategy in heart failure. Even though this would lead to inhibition of hypertrophy, there might be the danger of arrhythmia. On the contrary, targeted inhibition of *Gadlor1/2* effects in ECs and FBs could be beneficial, as it would entail increased angiogenesis and reduced FB-mediated fibrosis, which according to our data are likely the main underlying reasons for improved heart function in *Gadlor-*KO mice. Although more work is needed, we suggest that the inhibition of *Gadlor* effects in cardiac non-myocytes might be a promising strategy to treat heart failure in the future.

## Limitations

Although we identified CaMKII as direct target of *Gadlor1* and *2* in cardiomyocytes that crucially affect calcium cycling and hypertrophy, we did so far not identify the direct targets of *Gadlor* lncRNA in ECs or FBs, which will be our aim in future studies. Furthermore, we cannot prove the intercellular transport of *Gadlor1/2* in the heart *in vivo* at this point, as the detection of EV transfer of specific content between cells in the whole organ is methodologically impossible. While we have identified a crucial role of *Gadlor1/2* under pathological conditions, their role in normal physiology remains unclear. We envision *Gadlor1/2* might serve to fine-tune the link between capillary and cardiomyocyte growth and function also in the absence of pathology in the adult heart, but this needs experimental validation in the future.

## Acknowledgements

The authors thank the Core Facilities Pre-Clinical Models, Live Cell Imaging Mannheim (LIMA) and the Transgenic Mice Service Unit (TGSM) at Helmholtz Centre for Infection Research (HZI) for generating the knockout mice. The authors also acknowledge EMBL Proteomics Core Facility for performing proteomics analysis.

## Sources of Funding

This study was supported by grants from SFB1366 (CRC1366 - Vascular Control of Organ Function)/ 1-A6 (J.H.) by the Deutsche Forschungsgemeinschaft (DFG).

## Disclosures

The following patent has been granted to N.F., J.B. and J.H.: LncRNAs Gadlor1 and Gadlor2 for use in treating and preventing cardiac remodeling. US11208656 (granted 28.01.2021). The other authors declare that not no conflict of interest exists.

## Author contributions

**M.K** designed and performed the experiments, analyzed RNAseq data and wrote the first draft of the manuscript. **S.G** designed and performed the experiments and analyzed data. **N.F**, **F.A.T**, **R.W**, **S.H, S.L, G.M.D, M.S** and **R.H** performed experiments. **D.W** designed and supervised the generation of *Gadlor-*KO mice. **N.W** maintained the mouse colonies and performed experiments. **P.-S. K., Ja.H, D.F, S.U, K.B, A.M.G** and **J.C** provided experimental support for the study. **No.F., J.B., G.D.** and **T. W.** provide crucial advice for the study and critically revised the paper. **J.H.** designed and conceptualized the study, supervised all the experiments, and prepared the final draft of the manuscript. All authors read and approved the manuscript.

## Notes

### Competing Interest Statement

Competing Interest
The following patent has been granted to N.F., J.B. and J.H.: LncRNAs Gadlor1 and Gadlor2 for use in treating and preventing cardiac remodeling. US11208656 (granted 28.01.2021). The other authors declare that not no conflict of interest exists.

### Summary of Updates

The abstract of the manuscript was revised for misspelling and typos. There are no other changes were made in this version.

