## Supplemental Methods and Figures for "Endothelial cell-derived, secreted long non-coding RNAs *Gadlor1* and *Gadlor2* aggravate pathological cardiac remodeling via intercellular crosstalk"

### SUPPLEMENTAL MATERIAL

#### Extended Methods

#### Human Samples

Studies on human heart tissue samples were approved by the Institutional Ethical Board of Massachusetts General Hospital (US), where samples were collected from patients with end-stage heart failure undergoing cardiac transplantation. Control heart tissue samples were obtained from healthy organ volunteers when the organ was not eligible for transplantation, or from the victims of traffic accidents. Human serum samples were obtained from healthy blood donors, or from aortic stenosis patients before replacement of the aortic valve (clinical data are available in **Supplemental Table 1**), where all donors were provided a written informed consent for the collection and use of samples, that had received approval from the Institutional Review Board of Christian-Albrechts-Universität Kiel (File number: A174/09).

#### Measurement of GADLORs in Human Serum Samples

Human serum samples were collected on the day before Transcatheter Aortic Valve Implantation (TAVI). Serum-RNA was extracted using miRNeasy serum/plasma kit (#217184; Qiagen) based on the manufacturer's instructions. LncRNA levels were normalized by the addition of 2 µl of TATAA universal RNA Spike (#RS10SII; Tataa Biocenter) prior to RNA isolation. RNA was transcribed to cDNA (#K1652; Maxima H Minus First Strand cDNA Synthesis Kit; Thermo Fisher) and analyzed with TaqMan Non-coding RNA Technology (#4369016) using custom-specific TaqMan Gene Expression Assay (#43331348, ID: APPRKET und APRWEYP; Thermo Fisher). For quantitative RT-PCR, Stratagene's MX4000 multiplex qPCR system was used.

#### Animal Models and Studies

All studies including the use and care of animals were performed with the permission of the Regional Council Karlsruhe and the Lower Saxony State Office for Consumer Protection and Food Safety, Germany, with approved protocols 35-9185.81/G-144/18, I-22/03, 33.9-42502-12-10/0016, 33.19-42502-04-14/1403 and 33.8-42502-04-16/2356. Female and male animals were used in similar proportions throughout the study, except for the experiments in Figure 3, where only male mice were used.

Wildtype ICR/CD1 mice were in-bred in-house and C57BL/6N mice were obtained from Janvier Labs (Le Genest-Saint-Isle, France). Animals were maintained in a temperature-humidity controlled (22± 2°C and 35-60% humidity), 12-h dark-light cycled room with unlimited access to water and standard food.

Systemic *Gadlor* knock-out mice (*Gadlor*-KO) were generated by deletion of the respective region of mouse chromosome 16 using a CRISPR-Cas9-based strategy to achieve homozygous deletion. Briefly, embryonic stem cells (G4 line) were transfected with gRNAs LNC1-01 (target sequence: 5'-TTGTACATGAGCGGTTGTAG) and LNC2-01 (target sequence 5'-AGTATACAGGGGGTTACCAT) as well as in vitro transcribed CAS9-GFP mRNA. GFP-expressing cells were sorted and single-cell derived clones were established. 5 out of 72 analyzed clones showed extensive deletions between the gRNA target sites. Clones 1-37 comprised a deletion between 39,995,533 – 40,004,426 on one chromosome 16 allele. This clone was injected into C57BL/6 blastocysts to generate a transgenic mouse line which was then further crossed to achieve a homozygous deletion.

Pressure overload was induced in 8-10 weeks old mice by transverse aortic constriction (TAC) and maintained for 2 weeks for short-term and 8-12 weeks for long-term studies. Analgesia (0.1 mg/kg buprenorphine) was provided by subcutaneous injection. Tracheal intubation was followed by upper thoracotomy to visualize the aortic arch, which was tied with a 7-0 silk ligature around a 26-gauge needle that was removed immediately after secured constriction. Mice were injected with 0.1 mg/kg atropine, and the chest wall was closed with suture then followed by closure of the skin with surgical glue. The mice were ventilated with 2% isoflurane throughout the procedure. For postoperative care, additional analgesia was provided in drinking water for the following 5 days. To assess the mortality rate, mice were followed and inspected daily after TAC. Sham-operated animals were treated with the same procedure and medications, but no constriction was applied to the aortic arch. Mice were monitored with echocardiography before, and during pressure overload studies for assessment of cardiac function.

**Transthoracic Echocardiography**

Echocardiography was performed with 1% isoflurane anesthesia. Mice were dorsally placed on a heated table to maintain the body temperature during the procedure while the ECG and the respiration rate were recorded continuously. Echocardiography was recorded with a linear 30-40 MHz transducer (MX-550D) and the Vevo 3100 system (VisualSonics, Toronto, Canada). Data were analyzed with the Vevo Lab 5.5.0 software.

***Left Ventricle and Apical four Chamber View.*** Parameters were recorded in B-mode and M-mode in both parasternal long-axis (PSLAX) and short-axis (SAX) view at the papillary muscle level. Left-ventricular posterior wall thickness and left-ventricular end-diastolic volume were used to characterize LV microanatomy (LVPW and LVEDV) and the change of LV diameter length from end-diastole (LVID) to end-systole was used to assess contractility and to calculate LV ejection fraction and fractional shortening (LV-EF and LV-FS), which were measured with the PSLAX mode in the analysis. B-mode tracing in the apical four-chamber view was recorded at the atrioventricular valve level with pulse wave-Doppler (PW-Doppler) and tissue-Doppler measurements.

***Carotid flow measurement.*** To evaluate the degree of aortic constriction during TAC surgery, the peak velocity of flow in the right and left common carotid arteries (RCCA and LCCA) were measured 2 days after surgery with PW-Doppler at the level of carotid bifurcation to calculate the ratio of RCCA/LCCA flow.

### Organ Harvest

At the end of the experiment, mice were weighed and euthanized to collect the heart, lungs and liver. Organs were washed in cold PBS to remove the blood, and additionally, the heart was washed in 0.5% (w/v) KCl in PBS. The heart was transversely cut into half, and the upper layer was immediately embedded into OCT for histological analysis while the other half was snap frozen.

### Histology

OCT-embedded organs were sectioned into slices of 7 µm and 12 µm for immunofluorescence and Picro-Sirius red staining, respectively. Briefly, for immunofluorescence tissue sections were fixed with 4% PFA followed by permeabilization with 0.3% Triton-X in PBS and blocking with 3% BSA. Primary and secondary antibodies that were used are listed in **Supplemental Table 3**. Samples were visualized with Leica DMi8 Fluorescence microscope. Images were analyzed with Leica Application Suite X (LAS X) 3.7.

Sirius red staining was performed by fixing the tissue samples with 100% ice-cold acetone, and by following the standard protocol. Tissue sections were scanned with Zeiss Axio Scan.Z1. Images were analyzed with ZEN 2.6 Blue Edition, Carl Zeiss.

### Isolation and Culture of Juvenile Mouse Endothelial Cells (mECs)

Hearts from 7-12 days old mice were collected and washed in ice-cold DMEM high-glucose to clean and remove the atria. The minced tissue was transferred into 5 ml of enzyme solution (for 3-4 hearts) and incubated with collagenase I (500 U/ml, Worthington - LS004176) and DNase I (150 U/ml, Worthington - LS002139) in HBSS (without Ca and Mg). After digestion, the lysate was washed with FCS and passed through a 70 µm cell strainer. The washed lysate was incubated with CD31 antibody (BD, 553370) coupled to Dynabeads (Invitrogen, 11035) for the first step of isolation, before the cells were plated on 0.5% gelatinized plates. For culturing, cells were maintained for 3-4 days until they reached 80-90% confluency. During the first passage, further purification of cells was achieved by incubation with CD102 antibody (BD, 553370) coupled Dynabeads. Primary endothelial cells were maintained in DMEM with 20% FCS supplemented with non-essential amino acids and sodium pyruvate and passaged up to P3 for experiments.

**Isolation of Adult Mouse Endothelial Cells, Fibroblasts and Cardiomyocytes**

Isolation of endothelial cells and fibroblasts from adult mouse hearts was performed with MACS-magnetic beads from Miltenyi Biotec. Briefly, heart tissue was washed in ice-cold PBS and minced into small pieces before incubating in enzyme solution (collagenase I (500 U/ml, Worthington - LS004176) and DNase I (150 U/ml, Worthington - LS002139) in RPMI 1640 (Thermo Fischer Scientific: 31870025)). Digested tissue samples were passed through a 70 µm cell strainer and washed with FCS, and MACS buffer with BSA (Miltenyi Biotec, 130-091-222). After multiple washing steps, the tissue lysate was incubated with CD146 microbeads (Miltenyi Biotec, 130-092-007) before being passed through magnetic columns (Miltenyi Biotec, MS columns 130-042-201) for positive selection of endothelial cells (EC). The flow-through was incubated with feeder removal microbeads (Miltenyi Biotech, 130-095-531) and eluted after multiple washing steps to collect fibroblasts (FB). Isolated cells were stored as frozen cell pellet for later RNA or protein isolation for corresponding experiments.

Isolation of adult cardiac myocytes was achieved by using a Langendorff perfusion system. Briefly, immediately after excision, the heart was placed into ice-cold 1x perfusion buffer (10x stock: 1130 mM NaCl, 47 mM KCl, 6mM $KH_2PO_4$, 6 mM $Na_2HPO_4$, 12 mM $MgSO_4.7H_2O$, 120 mM $NaHCO_3$, 100 mM $KHCO_3$, 100 mM HEPES buffer solution and 300 mM Taurine) to clean the tissue around the aorta under a microscope. Then the aorta was placed onto a cannula (inner diameter 1mm) with forceps and tied with silk suture to allow perfusion through the coronary arteries. Subsequently, the heart was perfused with enzyme solution containing Liberase DH (5 mg/ml, Roche, 5401089001), trypsin (1%, Gibco, 15090046) and $CaCl_2$ (100 mM) to digest the tissue at 37°C. Enzymatic digestion was ended with Stop I and II solutions containing perfusion buffer with FCS and $CaCl_2$ (10 mM). Digested heart tissue samples were passed through a 100 µm cell strainer. Then, the calcium concentration was gradually increased by manual administration to digested heart tissue with continuous gentle mixing. Then the cells were subjected to either IonOptix analysis, culturing or direct RNA and protein extraction (after cardiomyocyte sedimentation).

**Isolation of Neonatal Rat Cardiomyocytes (NRCMs)**

Hearts from 1-3 days old rats were collected and washed with ice-cold 1x ADS (pH 7.35) to clean the blood and remove the atria. The ventricular parts were minced in 1x ADS and digested in enzyme solution containing collagenase Type II (Worthington: LS004176) and pancreatin (Sigma P3292). The resulting cells were loaded onto a Percoll gradient to separate cardiomyocytes (NRCMs) and non-cardiomyocytes. NRCMs were seeded ($4.0\times10^5$ /well for 6 well-plate) onto 0.5% Gelatine-coated plates.

**Isolation of Extracellular Vesicles (EVs)**

Isolation of EVs was performed with ultracentrifugation from the supernatant of C166 cells that were cultured with EV-depleted FCS. Initially, cell debris and large vesicles (apoptotic bodies) were removed by centrifugations at 300xg and 10.000xg, respectively. Subsequently, small EVs (including microvesicles and exosomes) were collected with 100.000xg ultracentrifugation for 90 minutes, and similarly washed with PBS, before a second round of centrifugation was performed. The EV pellet was re-suspended in PBS for NTA (ZetaView Nanoparticle Tracing Videomicroscope, PMX-120) measurements, or with QIAzol lysis buffer (Qiagen, 79306) for RNA isolation.

***EV-RNA isolation.*** RNA isolation from EVs was performed by RNA precipitation as described before. Briefly, the EV pellet was resuspended with 1 ml of QIAzol, and then mixed with 200 µl of chloroform for phase separation. Samples were mixed thoroughly and incubated for 5 minutes at room temperature (RT), then centrifuged at 12.000xg at 4°C for 15 minutes. The aqueous upper layer was mixed with 10% (v/v) sodium acetate (3M, pH 5.5) and 4 µl of glycogen (5 mg/ml) in 1.5 ml ethanol (100%). Samples were incubated at -80°C overnight, then centrifuged at 16.000xg for 30 minutes to pellet the RNA. The RNA pellet was then washed with 70% ethanol and dried, before being re-suspended in nuclease-free water.

***EV RNase and Proteinase-K Protection Assay.*** To test the transfer of *Gadlor* lncRNAs within EVs, we performed RNase and proteinase K protection assays with and without prior the application of Triton-X. In brief, EVs were isolated and treated with 100 ng/µl of RNase A, and 20 mg/µl of proteinase K for 30 minutes at 37°C. 5 mM of PMSF was used to inactivate the proteinase K at RT. To disrupt the lipid bilayer, EVs were incubated with 1% Triton-X for 1 hour for selected samples. As a negative control, RNA isolated from EVs was also treated with the same protocol.

***EV-labelling with PKH67 cell linker.*** To visualize EVs for transfer and re-uptake experiments *in vitro,* they were labelled with PHK67 Green Fluorescent Cell Linker kit (Sigma MIDI67). Briefly, the EV pellet was re-suspended with 1 ml of diluent C including 4 µl of PKH67 dye, and incubated for 5 minutes at RT. Then, 1% BSA-PBS was added into the suspension to remove the unspecific binding of the dye. Then EVs were pelleted at 3000xg for 1 hour, which were then suspended with the cell culture medium and added on NRCMs. Visualization of EVs was conducted with the Leica Confocal Microscope TCS SP8 and images were analyzed by Leica Application Suite X (LAS X) 3.7.

***EV FACS Staining.*** Characterization of surface markers of EVs was performed with flow cytometry by staining EC-derived EVs with CD9-PerCP-Cy5.5 (Miltenyi Biotec, 130-102-278, Clone: MZ3), CD63-APC (Miltenyi Biotec, 130-108-894, Clone: REA563) and CD54-FITC (BD, 553252, Clone: 3E2) labelled antibodies for 1 hour at 4°C. Measurements were obtained with BD FACSCanto II.

***EV Transmission Electron Microscopy (TEM).*** Visualization of EVs was performed with TEM. After centrifugation, pellets were resuspended in the minimum possible volume of residual liquid and 25 % aqueous glutaraldehyde was added to a final concentration of 1 %. After fixation overnight, samples were mixed 1:1 (v/v) with 4 % agar at 40 °C. After hardening, the agar blocks were cut into cubes of 1 mm in size. Further preparation of samples and electron microscopy were performed as described, before being imaged with a FEI Morgagni 268 transmission electron microscope (FEI, Eindhoven, Netherlands) operated at 80 kV using a Veleta CCD camera (Olympus Soft Imaging Solutions).

***EV-mediated Gadlor1 and Gadlor2 overexpression.*** As a gain-of-function approach, *Gadlor1* and *Gadlor2* were overexpressed in mouse hearts by administration of *Gadlor1/2*-enriched EVs. To this end, C166 endothelial cells were infected with *Gadlor1* and *Gadlor2* adenoviruses for 48 hours. EVs were collected with ExoQuick-TC solution (EXOTC50A-1,65 SystemsBio) based on manufacturers' protocols with some adjustments. In vivo overexpression of *Gadlor* lncRNAs was achieved by injecting EVs directly into the left ventricles before TAC surgery. The intra-ventricular injection was performed by simultaneous cross-clamping of the aorta and pulmonary artery distal of the origin of the coronary vessels.

**Cell Culture**

C166 (CRL-2581, ATCC) mouse embryonic endothelial cells and NIH3T3 (CRL-1658, ATCC) mouse fibroblast cells were cultured in Dulbecco's Modified Eagle's Medium (DMEM) with 10% FCS, while MCECs (CLU510-P, Tebu-bio), immortalized mouse cardiac endothelial cells, were cultured with 5% FCS containing DMEM supplemented with L-glutamine (1%), penicillin/ streptomycin (1%) and HEPES. HL-1 cardiac muscle cells (SCC065, Merck) were cultured with Claycomb media (Sigma, 51800C) supplemented with 10% FCS, 0.1 mM norepinephrine, 2 mM L-glutamine and 1X penicillin-streptomycin (P/S). HL-1 cells were plated on 0.1% Gelatine-coated flasks, and cell culture media was refreshed daily. Cells were cultured for up to 10 passages.

***Adenoviral infection.*** For overexpression of *Gadlor* lncRNAs, recombinant adenoviruses were used. Mouse cDNAs of AK037972 (*Gadlor*1) and AK038629 (*Gadlor*2) were subcloned into the pShuttleCMV vector (Source Bioscience, UK) and adenoviruses were generated by the AdEasy Adenoviral Vector system (Agilent, 240009). An adenovirus overexpressing β-Galactosidase (Adß*gal*) was used as a control. After the production of adenoviruses, the viral titer was calculated with the help of the AdEasy Viral Titer Kit (Agilent, 972500), and all the experiments were performed with 50 MOI. Adenoviral infection on cultured cells was performed for 4 hours at 37°C on cells with media containing heat-inactivated FCS.

***Co-culture experiments.*** For co-culture experiments, adult mouse cardiomyocytes isolated from *Gadlor*-KO mice were seeded on laminin (Santa Cruz, SC-29012) coated 6-well plates and cultured with MCECs plated on 1 µm pore-sized inserts (Thincert, Greiner, 657610). Cells were collected separately after 48 hours for RNA isolation.

To inhibit EV trafficking between MCECs and CMs in the co-culture model, MCECs were pre-treated with 20 µM of GW4869 (Hoelzel Biotech) for 6 hours, and then placed for 48 hours of co-culture with isolated *Gadlor*-KO or WT CMs.

**Endothelial Cell Sprouting Assay**

To study the effect of *Gadlor* lncRNAs on angiogenesis, we performed sprouting assays with C166 mouse ECs according to a well-established protocol previously described. Briefly, cells were infected with *Ad. ßgal or Ad.Gadlor1/2* adenovirus for 24 hours. 50.000 cells were re-suspended with 4 mL DMEM + 10 % FCS and 1 mL methylcellulose solution (Sigma-Aldrich, M0512) and drops containing 25 µl of suspension were pipetted onto a 10 cm cell culture dish. The drops were then incubated upside-down in a cell culture incubator for 24 hours to allow spheroid formation. On the following day, the cell spheroids were gently washed off the hanging drops with PBS, collected by 200xg centrifugation for 5 minutes and re-suspended with methylcellulose solution containing 20 % FCS. The collagen matrix was prepared in parallel on ice with collagen stock solution (Corning, 11563550), diluted in 10x M199 medium (Sigma-Aldrich, M0650) and the required amount of sodium hydroxide to set the pH for polymerization. Then the collagen medium was mixed with the spheroids. 1 mL of the spheroid-collagen-methylcellulose mixtures were added per well in a 24-well plate and incubated in a cell culture incubator for 30 min to induce the polymerization of the collagen-methylcellulose matrix. The endothelial spheroids were stimulated with 100 µl of DMEM + 10 % FCS, 25 ng/mL FGF2 or 10 ng/mL TGF-β1 by adding it dropwise to the collagen matrix. After 24 hours, the sprouting assay was stopped by adding 1 mL of 4% paraformaldehyde to the wells. Spheroids were visualized with the 10X objective in bright-field microscopy and analyzed with ImageJ.

**RNA Isolation and qRT-PCR**

RNA isolation from tissue samples was performed with QIAzol reagent (Qiagen, 79306), and from isolated cells with NucleoSpin RNA isolation kit (Macherey-Nagel, 740955.250) according to manufacturers' protocols. Samples were incubated with rDNase (Macherey-Nagel, 740963) on the silica-columns to remove any DNA contamination. cDNA was generated by using the Maxima H minus First strand cDNA synthesis kit (Thermo Fisher Scientific, K1652). Quantitative PCR was performed with the Maxima SYBR Green mix with ROX as reference dye (Thermo Scientific, K0253) on an AriaMx Real-time PCR System (Agilent, G8830a). Gene expression was normalized to *Gapdh*,18S or *U6* expression. All qPCR primer sequences are listed in **Supplemental Table 4**.

**Protein Isolation and Western blotting**

Heart tissue protein lysates were prepared from frozen pulverized tissue with lysis buffer containing 30 mM Tris-pH 8.8, 5 mM EDTA-pH 8.0, 3% SDS (v/v), 10% glycerol (m/v), protease and phosphatase inhibitors. Protein quantification was performed by BCA assay (ThermoFisher, Pierce BCA Protein Assay kit, 23225). Samples were incubated at 95°C for 5 minutes after the addition of Laemmli buffer (1X final concentration) and proteins were separated with SDS-PAGE electrophoresis. The following primary antibodies were used: Calsequestrin (Thermo Scientific PA1-913, rabbit), Phospholamban, pThr17 (Badrilla A010-13, rabbit), Phospholamban (Badrilla, A010-14 mouse). The following secondary antibodies were used: Rabbit anti-mouse peroxidase (Sigma-Aldrich), and goat anti-rabbit peroxidase (Sigma-Aldrich).

**Sarcomere Contractility and Calcium Transient Measurements**

Adult cardiomyocytes were isolated 1-week after TAC operation. Cardiomyocytes were plated on laminin coated (10 µg/cm$^2$) 35 mm dishes (MatTek 10 mm Glass bottom dishes, P35G-1.5-10-C) and gently washed 1 hour later with MEM medium without butanedione monoxime (BDM). The plated cells were then transferred into the Ion-Optix Multicell High-throughput System chamber, where they were simultaneously paced (18 V, 2.5 Hz, 4 ms impulse duration), and assessed for sarcomere length and contraction. Calcium handling was recorded after incubation with 1 µM fura-2, AM (Invitrogen, F1221).

For measurements following *Gadlor1/2* overexpression via EV-mediated transfer, isolated adult cardiomyocytes were incubated with EVs for 4 hours. To assess the effect of CaMKII inhibition, cells were treated with KN93 (1 µM) for 1 hour before the measurements. The recordings were analyzed with IonWizard software.

**RNA antisense purification coupled with mass spectrometry (RAP-MS)**

To identify the protein interaction partners of *Gadlor* lncRNAs we performed RAP-MS as described previously with minor modifications.[53] The 5' biotinylated antisense *Gadlor1* and *Gadlor2* probes were pooled in the experiment and the sequences are listed in **Supplemental Table 5**. We used HL-1 cardiomyocytes overexpressing *Gadlor1* and *Gadlor2* after infection with Ad.*Gadlor1* and Ad.*Gadlor2.* RNA antisense purification (RAP) was performed with cells collected from five 15 cm dishes per sample, where each experimental group contained three replicates without (as a negative control to identify non-specific 'background' proteins) and four replicates with UV cross-linked conditions. HL-1 cells were washed with cold PBS twice and cross-linked using 150 mJ/cm$^2$ of 254 nm UV light. Cells were then lysed with lysis buffer, incubated for 10 minutes on ice, homogenized by passing several times through a 21G needle and DNA was digested with addition of DNase salt solution and Turbo DNase for 10 minutes at 37ºC (ThermoFisher, AM2238). Hybridization conditions were adjusted by the addition of an equal amount of hybridization buffer. Lysates were precleared with streptavidin-coated magnetic beads. Biotin-labelled *Gadlor1* and *Gadlor2* probes were heated to 85ºC for three minutes and then incubated with the lysate for 2 hours at 67°C. Probe-RNA complexes were captured by pre-washed streptavidin-coated magnetic beads and incubated at 37ºC for 30 minutes. Lysate was removed from beads by magnetic separation, and beads were washed four times in hybridization buffer at 67ºC. *Gadlor* lncRNA-bound proteins were then released by Benzonase RNA digestion for two hours at 37ºC. The captured protein samples were identified by TMT labelling followed by liquid chromatography-mass spectroscopy (LC-MS/MS) by the EMBL proteomics core facility.

**RNA immunoprecipitation (RIP)**

We performed RIP with minor modifications to a previously described protocol to confirm the interaction of *Gadlor1* and *Gadlor2* lncRNAs with CaMKII. HL-1 cardiomyocytes were used to overexpress *Gadlor* lncRNAs by adenovirus treatment, while Ad.βgal treated samples were used as control. Three 15 cm plates were combined for preparation of each sample. Briefly, cells were washed with ice-cold PBS twice and lysed with polysome lysis buffer (100 mM KCl, 5 mM MgCl2, 10 mM HEPES pH 7, 0.5% IGEPAL CA-630, 0.1 mM DTT, 1 x protease inhibitor cocktail, 1 x RNasIN). Cell lysates were passed through a 26G needle multiple times for homogenization. Protein G magnetic beads (Bio-Rad SureBeads, 161-4221) were washed twice with NT-2 buffer (50 mM Tris-HCl pH 7.4, 150 mM NaCl, 5 mM MgCl2, 0.05 % IGEPAL CA-630) and coupled with anti-CaMKII antibody (Santa Cruz, sc-5306) or anti-IgG (mouse, Cell Signaling 7076, as control) overnight at 4°C. Antibody coupled beads were washed twice with NT-2 buffer and resuspended in NET-2 buffer (50 mM Tris-HCl pH 7.4, 150 mM NaCl, 5 mM MgCl2, 0.05 % IGEPAL CA-630, 20 mM EDTA pH 8, 1 mM DTT, 1 x RNaseIN). Cell lysates were added to beads and incubated at 4°C for 2 hours (10% lysate was removed as input and stored on ice prior to IP).

After supernatant was removed from the beads, 1 ml of ice-cold NT-2 buffer was used to wash the beads five times in total. Then, beads were resuspended with Proteinase K buffer (1x NT-2 buffer and 1% SDS) and incubated at 55°C for 30 minutes with constant shaking while input sample conditions were adjusted accordingly and processed in parallel. The supernatant was transferred into fresh tubes and combined with NT-2 buffer and UltraPure phenol-chloroform (ThermoFisher, 15593031) for RNA purification. After centrifugation in heavy-lock tubes at 15.000 x g for 15 minutes, RNA was purified from the clear aqueous phase with the RNA Clean and concentrator kit (Zymo Research, R1017).

**RNAScope *in situ* Hybridization**

In situ hybridization was performed to localize Gadlor1 (Mm-LOC118567341-C3), Gadlor2 (Mm-AK038629-C1, Cat no: 404391) and Cdh5 (Mm-Cdh5-C2, Cat no: 312531) mRNA expression on mouse heart tissue slides with RNAscope Multiplex Fluorescent v2 kit. The assay was performed according to the manufacturer’s protocol. Briefly, OCT-embedded tissue slides (7 µm thickness) were used in the assay, fixed with 4% PFA and then stepwise dehydration was performed with increasing concentrations of ethanol. Tissue slides were subsequently treated with hydrogen peroxide (10 min at RT) and protease IV (30 min at RT) in the humidity control incubator. Slides were incubated for 2 hours with the mixture of probes to hybridize with the corresponding RNAs. Next, the signal was amplified with RNAScope Multiplex FL v2 AMP 1-3 reagents and then HRP channels were developed sequentially. Finally, DAPI was used to counterstain the slides and each was mounted with ProLong Gold Antifade Mountant (Fisher Scientific).

**DNA Isolation and Analysis of mtDNA/ nDNA**

To analyze the mitochondrial DNA (mtDNA) to nuclear DNA (nDNA) ratio of *Gadlor*-KO and WT mouse hearts after 2-weeks of TAC, genomic DNA was isolated from mouse heart tissue samples obtained after TAC (2w) with a DNA isolation kit according to manufacturers' instructions (PureLink Genomic DNA Mini kit, Invitrogen, K182001). The mtDNA/ nDNA ratios were quantified by quantitative PCR using ND1 and 16S rRNA primers listed in **Supplemental Table 4**.

**Bulk RNA sequencing and Bioinformatics**

To perform RNA sequencing, RNA isolation was performed from isolated cells as described at indicated time points (2 weeks after TAC or sham surgery). Quality control of RNA samples (Agilent 2100 Fragment Analyzer), and library preparation (DNBSEQ Eukaryotic Strand-specific mRNA library) were performed by BGI, Hong Kong. Bulk RNA sequencing from different cardiac cells were performed as stranded and single-end with 50 base sequence read length by BGI with Illumina HiSeq 2500. Initially, the trimming of adapter sequences from fastq files was performed with the R package *FastqCleaner*. For aligning the reads to the reference genome (mm10) after trimming, the R package *bowtie2* alignment tool was used. Gene annotation was performed with the *bioMaRt* R package. Library size of the samples was normalized to counts per million (cpm) and transformed into log2 values followed by calculation of differential gene expression with the *edgeR* package of R. Significant changes in gene expression between compared groups were filtered based on FDR < 0.05, and fold change (FC) > 1.5. Gene ontology analysis were performed with Metascape and DAVID online tools. Heatmaps that show the differentially regulated genes were generated by heatmap.2 function in *ggplot2* library in R.

**Data Availability**

The authors declare that the data supporting the findings of this study are available within the paper and its supplementary information. RNA sequencing data sets were deposited in National Center for Biotechnology Information's (NCBI) Gene Expression Omnibus (GEO) repository with the accession number GSE213612.

**Statistics**

Data analysis and statistical analysis were performed with GraphPad Prism software (Version 8). Data are shown as mean ± standard error of the mean (SEM). All the experiments were carried out in at least 3 biological replicates. The number of replicates for animal experiments and cell culture experiments are indicated in the figure legends. The investigators were blinded for mouse genotype and treatment during surgeries, echocardiography, organ weight determination and all histological and immunofluorescence quantifications. Premature death was a criterion for exclusion from an ongoing two weeks TAC experiment. Death rates were not significantly different between experimental two weeks TAC groups.

Initially, all the datasets were analyzed for normality to allow the application of the proper statistical test. An unpaired 2-tailed Student t-test was used for comparing 2 groups only. Comparing multiple groups for one condition was performed with one-way ANOVA and Fisher's LSD post-hoc test. Comparing multiple groups for multiple conditions was achieved with two-way ANOVA and followed with Fisher's LSD post-hoc test when applicable. For non-parametric data sets, the Mann-Whitney and Kruksal-Willis tests were used to compare two groups and multiple groups, respectively. Values of $p<0.05$ were considered statistically significant.

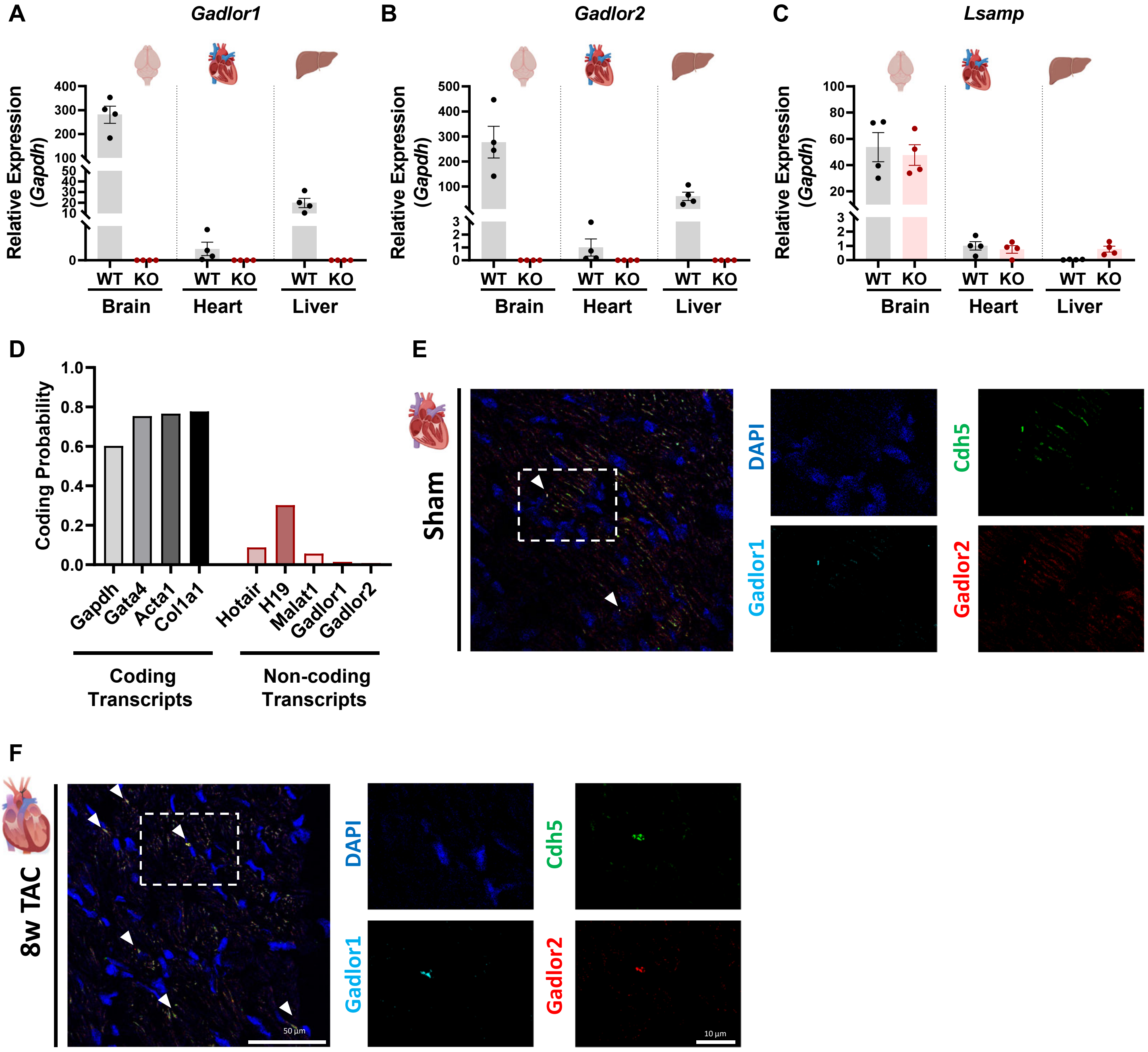


**Supplemental Figure 1: *Gadlor1* and *Gadlor2* are non-coding transcripts that are located in the intronic region of the *Lsamp* gene.**

**A-C.** Systemic deletion of the whole region containing *Gadlor* lncRNAs did not affect the expression of the neighbor gene *Lsamp,* which is mainly expressed in brain tissue. (*Gapdh* expression used for normalization) Relative expression of *Gadlor1, Gadlor2* and *Lsamp* were shown in mouse brain, heart and liver tissue samples collected from WT and *Gadlor*-KO mice (n=4, same samples were indicated in each graph). **D.** Coding probability of *Gadlor1* and *Gadlor2* was evaluated with the online Coding Potential Assessment Tool (CPAT - http://lilab.research.bcm.edu/) and compared with coding transcripts and well-known lncRNAs for validation that revealed *Gadlor1* and *Gadlor2* are non-coding transcripts. **E-F.** Fluorescent *in situ* hybridization of *Gadlor1* (cyan), *Gadlor2* (red) and *Cdh5 RNAs* in mouse heart tissue sections of **E.** sham and following **F.** 8-weeks of TAC surgery. *Cdh5* shows endothelial cells (green), nuclei stained with DAPI (blue), and the overlap in 3 channels was indicated with arrows. Dashed rectangles were the zoomed areas showed in separate pictures. Scale bars: 50 $\mu$m and 10 $\mu$m, as indicated.

**A** **AK037972 *(Gadlor1)***

```
#=======================================
# Aligned_sequences: 2
# Gap_penalty: 10.0
# Extend_penalty: 0.5
#
# Length: 578
# Identity:      268/578 (46.4%)
# Similarity:    268/578 (46.4%)
# Gaps:          259/578 (44.8%)
# Score: 740.5
#=======================================

H   1 AAATTGTCTAGAA-TGTGGCAGTTTAAAAACACATCTACATTGATCTACA     49
      |||||||.||||| |||||||.|||||||.|||.||..|||||||||||.
M   1 AAATTGTTTAGAACTGTGGCACTTTAAAAGCACCTCGGCATTGATCTACT     50

     50 GTCTCAAGGTCCCAC--CTTTGTGTTACTTTTAGAATTTGTAGATGGATG     97
        ..|||.||.||.|||  ||.||||.||||||||||.|||||||||||||.
     51 AACTCTAGATCTCACCTCTCTGTGCTACTTTTAGAGTTTGTAGATGGATA    100

     98 TTATTCAGCA-AGGATGACTCATCTACATT-TGGTTTGTCAATTTGTTTT    145
        .|.|.||||| |||| ||.||||||||.|| ||  .||||.|||||||||
    101 CTGTCCAGCAGAGGA-GAGTCATCTACGTTCTG--ATGTCGATTTGTTTT    147

    146 CAAAGATCATA----TGTACATAGCGTCCT----TCGATCTTACTTTCCC    187
        ||.||||||.|    || ||||    ||||    |.|     |||.||||
    148 CAGAGATCAGACACTTG-ACAT----TCCTAGAGTGG-----ACTGTCCC    187

    188 TTTAAAATATGAAACTGATAATGACCTTGAAAAAGAT-------AGAATT    230
        |||||||||||||||...|||||.|||..|.||||||       |.|.||
    188 TTTAAAATATGAAACAAGTAATGGCCTACAGAAAGATTATGATAAAACTT    237

    231 TC----------------------------------------AG------    234
        ||                                        ||
    238 TCCAGTTTTTTGGCTTTTCCCTTCGGCCCTGACTGGATGTGTAGTCTACT    287

    235 --CCACAA------------------------------------------    240
          ||||||
    288 TCCCACAATCCTGTATGTTACATGGTTCCCTACCCAGACCACTCCCTTTG    337

    241 -----------------------------------------GCATCCCA-    248
                                                 ||||.|||
    338 CCCCACCCCAAAAATGGCTACTGGGCTTTCTCAATCATCTTGCATGCCAC    387

    249 ---------------------------------TGAA-------AT----    254
                                         ||||       ||
    388 CCCTCGCCTTCTTTTATGTTTCAGAGTTGTGTGAAGAATCACATTAGT    437

    255 --TAATGTATCCATACTTTTAACA--CTA-----GTATAC-----TTTAT    290
          ||.|||||.|.|.||||| |||  |||     ||.|||     |||||
    438 GATACTGTATGCTTTCTTTT-ACAATCTATTTACGTTTACACATATTTAT    486

    291 CTTCTTATTT-CA-AATGCTC--TGTCTCC--------------------    316
        .||..||||| || |.|.|||  |||.|.|
    487 TTTTATATTTACATATTTCTCATTGTTTACCAGTTGGCAATAGTGATATT    536

    317 TT---------CTGTCCA--TTACTTCC    333
        ||         ||.|.||  ||||||||
    537 TTGTGAGCTGACTTTACATTTTACTTCC    564
```

**B** **AK038629 (*Gadlor2*)**

```
#=======================================
# Aligned_sequences: 2
# Gap_penalty: 10.0
# Extend_penalty: 0.5
#
# Length: 913
# Identity:      656/913 (71.9%)
# Similarity:    656/913 (71.9%)
# Gaps:          130/913 (14.2%)
# Score: 2336.5
#=======================================

H   1 AAAAAGAAAATTTGGA-AGATGCTCAAATAGGA-TTGATAGAGGTTCATT     48
      ||.||.|||||||||| || |||.||.|||||| ||| .||.|..|||..
M  34 AAGAATAAAATTTGGACAG-TGCACACATAGGACTTG-CAGGGACTCACA     81

     49 TTGTGTCATATTTT-CGCTTTATATGGATGTTGATATGTTCTCAGAATAA     97
        .||||||||||||| |.||||.||||.|||||||||||||.||||||| |
     82 GTGTGTCATATTTTGCACTTTTTATGAATGTTGATATGTTTTCAGAAT-A    130

     98 ATTTATAACTGCAATAGAAAATGGAATGACTATTGATTATTTTTAGTTA    147
        |||||||||||||||.|||||..|||||.||||||||||..||||||||.
    131 ATTTATAACTGCAATTGAAAACTGAATGGCTATTGATTAAATTTTAGTTT    180

    148 CCCAAACTAATTGTCATATAAAGCTTTGTAAAT-CTTTTA-AAATGAGTA    195
        |..||||||||||.||||||||||||..||||| |||||| |||  ||.|
    181 CAAAAACTAATTGACATATAAAGCTTCATAAATCCTTTTAGAAA--AGGA    228

    196 TGGGTGTAAATTAAAAACGAATTAAAGCAGAGGCTTATAAAGACACT---    242
        ||.||.||||||||||||.|||||.||||.||||||||||| |.|||
    229 TGAGTATAAATTAAAAACTAATTACAGCAAAGGCTTATAAA-ATACTCCA    277

    243 ATCTCCCCTTCAATGGGTTATGTATTTTTGTTGTGTGTTAGGAAGATATC    292
        || |||..|||.|| ||||||.||   .||||||.||.||.||||.|.|.
    278 AT-TCCAGTTCTAT-GGTTATATA---CTGTTGTATGCTATGAAGGTGTT    322

    293 TCTTGATCTCCACACCCAAAACTCCTTCC----------AAATGAACAAA    332
        .|||..|||.|..|..||||||||||.||          |.||||||
    323 CCTTTGTCTTCTTAATCAAAACTCCTGCCATAGAATTAAATATGAAC---    369

    333 GGCCTACACACAAAAATTATTTCAATTGCTAATAGCCAGTTTTTAT----    378
                ||.||||..|||||.||||||||||....|.|||.|||
    370 --------CATAAAAGATATTTTAATTGCTAATGATTACTTTATATATAG    411

    379 ------TTTTCAAAAGTAAATGCTTTCTTTTTTTGAAAAGTAAATATAAC    422
              |||||||.||                      |||  .|||||.
    412 TATCCCTTTTCAAGAG----------------------AGT--TTATAAG    437

    423 TATTTTAACAAAGTAAGATTTAAAAA---AAAAAACAACTCAGAAAAAAG    469
        .|||||.|...|||||.|||||||||   ||||||.|||      |.|||
    438 AATTTTTAATCAGTAATATTTAAAAATTTAAAAAAGAAC------AGAAG    481

    470 T--GCCAT------GCAG-AGATAACAGGGCTGATCTGTCTGTTATCTGC    510
        |  |||||      |||| ||| |.||      || ||.|.||||||||.
    482 TGGGCCATGTTAAGGCAGCAGA-ACCA------AT-TGCCAGTTATCTGT    523

    511 AGCCTCTATCCTTACTGTTAACAAGCCTTTTATCTTTGAAGACACTAAAC    560
        |.|||||||||||..||.|||.||||||||||.|||.|||||||||||.|
    524 AACCTCTATCCTTGATGCTAATAAGCCTTTTACCTTCGAAGACACTAAGC    573

    561 ATCTGGGATCTAAGCACTGACACCTATTAGTTACAGTGGTTTCCTTTTAC    610
        ||||||||||||||.|.||||||||||||||||||.||||||||||||||
    574 ATCTGGGATCTAAGTAGTGACACCTATTAGTTACAATGGTTTCCTTTTAC    623

    611 CTTTCTAAACTATCTGATAGATAAACCTCCAGGAATCCTACAAAATTAGG    660
        | ||||||.|.|||||||||||||||||||||||||||.||||||||.|.
    624 C-TTCTAAGCAATCTGATAGATAAACCTCCAGGAATCCCACAAAATTGGA    672

    661 CCCTTAATTGACCAAACGAATGTCTGTCTCTCTTTTTCAATCTAAATCAA    710
        ||||||||||||||||.|||||.|||||||||||||.||.|.|||||..|
    673 CCCTTAATTGACCAAATGAATGGCTGTCTCTCTTTTACAGTATAAATTGA    722

    711 AAAG-GAGTTTGCTCCTAGGAG--ATGAGATGATGTACAGTGGAAAAAAA    757
        |||| |||||||||||.|.|||  |.||.||.|..||    |||||.|||
    723 AAAGTGAGTTTGCTCCCATGAGGTAAGACATAACTTA----GGAAAGAAA    768

    758 TATTTCA-GCTGCAACTCCACTGGTAACCGTCAATGTGACCTTATAATCT    806
            ||| ||||..||||||||.|..| |.|.|.||||.||||.||||||
    769 ----TCACGCTGAGACTCCACTTGCCA-CATTAGTGTGCCCTTGTAATCT    813

    807 CCCC-----CCAATTTATTCCCA-AAACAGCTA--CCAAAGTGATCTTTC    848
        |.||     |||       |||| .||||| ||  .|||||||||..|||
    814 CACCAAAAGCCA-------CCCACCAACAG-TAGGACAAAGTGATTCTTC    855

    849 TAAAACCCACATC    861
        ||||||.||||||
    856 TAAAACGCACATC    868
```

**C**

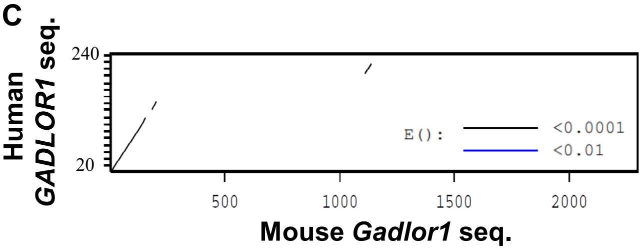


**D**

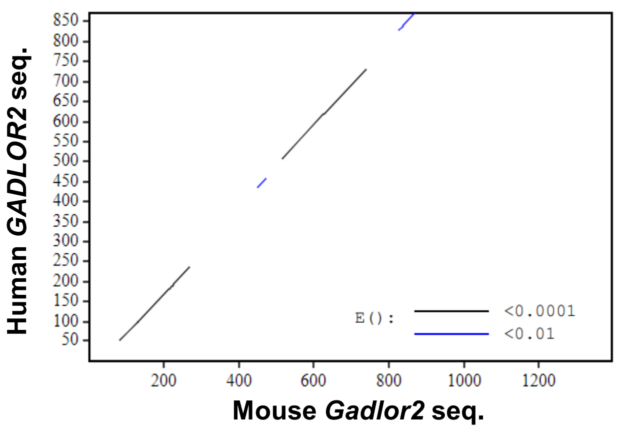


**Supplemental Figure 2: Conservation levels of *Gadlor1/2* between mouse and human genome.**

**A-B.** Assessment between human (H) and mouse (M) *Gadlor1/2* in sequence conservation level evaluated with EMBOSS-Water local alignment tool (https://www.ebi.ac.uk/Tools/psa/emboss_water/), and **C-D.** visualization of pairwise local alignment with LALIGN DNA:DNA tool of University of Virginia (https://fastademo.bioch.virginia.edu/fasta_www2/fasta_www.cgi?rm=lplalign).

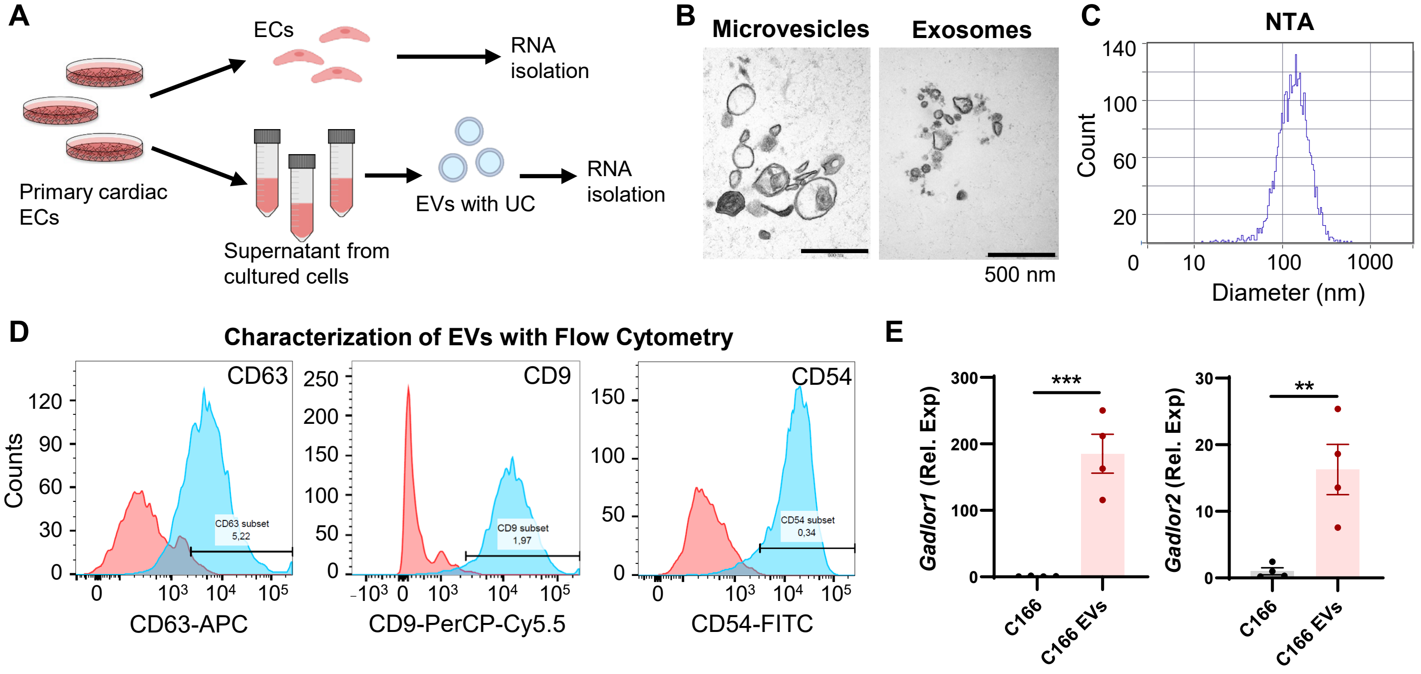


**Supplemental Figure 3:** ***Gadlor*** **lncRNAs are mainly secreted in endothelial cells (EC) derived extracellular vesicles (EVs).**

**A.** Scheme depicting the experimental design of EV isolation with ultracentrifugation (UC) from cultured primary cardiac ECs. Characterization of cardiac EC-derived EVs by **B.** visualizing microvesicles and exosomes with electron microscopy (scale bar: 500 nm) and **C.** detecting the size in diameter (nm: nanometers) with Nanoparticle Tracking Analysis (NTA). **D.** Flow cytometry analysis of EV surface markers (CD63-APC, CD9-PerCP-Cy5.5 and CD54-FITC) on EC-derived EVs that were stained against markers (blue curve) and compared to isotype control (red curve). **E.** Expression of *Gadlor1* and *Gadlor2* in C166 mouse endothelial cell and in EVs derived from these cells. Data are shown as mean±SEM. Data normality was evaluated with Shapiro-Wilk test and p-values were calculated with Student's t-test for parametric (or Mann-Whitney for non-parametric) assessment. *p-value<0.05, **p-value<0.01, ***p-value<0.001.

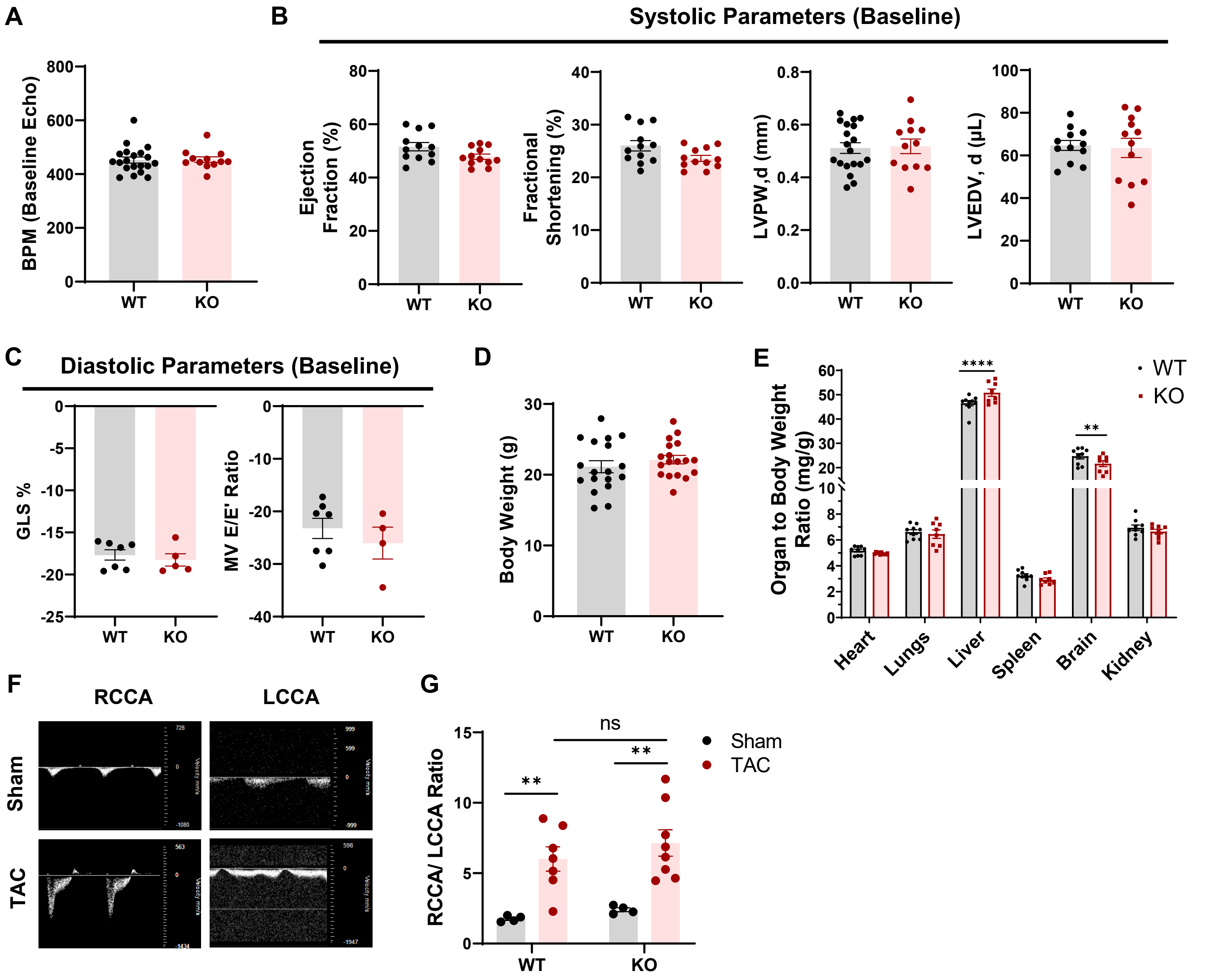


**Supplemental Figure 4: Baseline phenotypic characterization of *Gadlor*-KO mice compared to wild-type (WT) littermates.**

**A.** Heart rate of WT and *Gadlor*-KO mice during echocardiography (BPM: beats per minute, n≥12). **B.** Analysis of systolic function parameters of left ventricle (LV) ejection fraction (%), fractional shortening (%), LV posterior wall thickness in diastole (mm) and LV end diastolic volume (µl) (n≥12). **C.** Analysis of diastolic function parameters, global longitudinal strain (GLS, %) and mitral valve (MV) E to E' ratio (n≥4). **D.** Body weight (g) of adult (9 weeks old) WT and *Gadlor*-KO animals (n≥18), and **E.** organ to body weight ratio (mg/g) of isolated organs including heart, lungs, liver, spleen, brain and kidney (n≥8). **F-G.** Representative images and quantification of flow measurement in right and left common carotid arteries (RCCA and LCCA) in sham (n≥4) and TAC (n≥7) mice to measure the strength of aortic constriction in WT and *Gadlor*-KO animals.

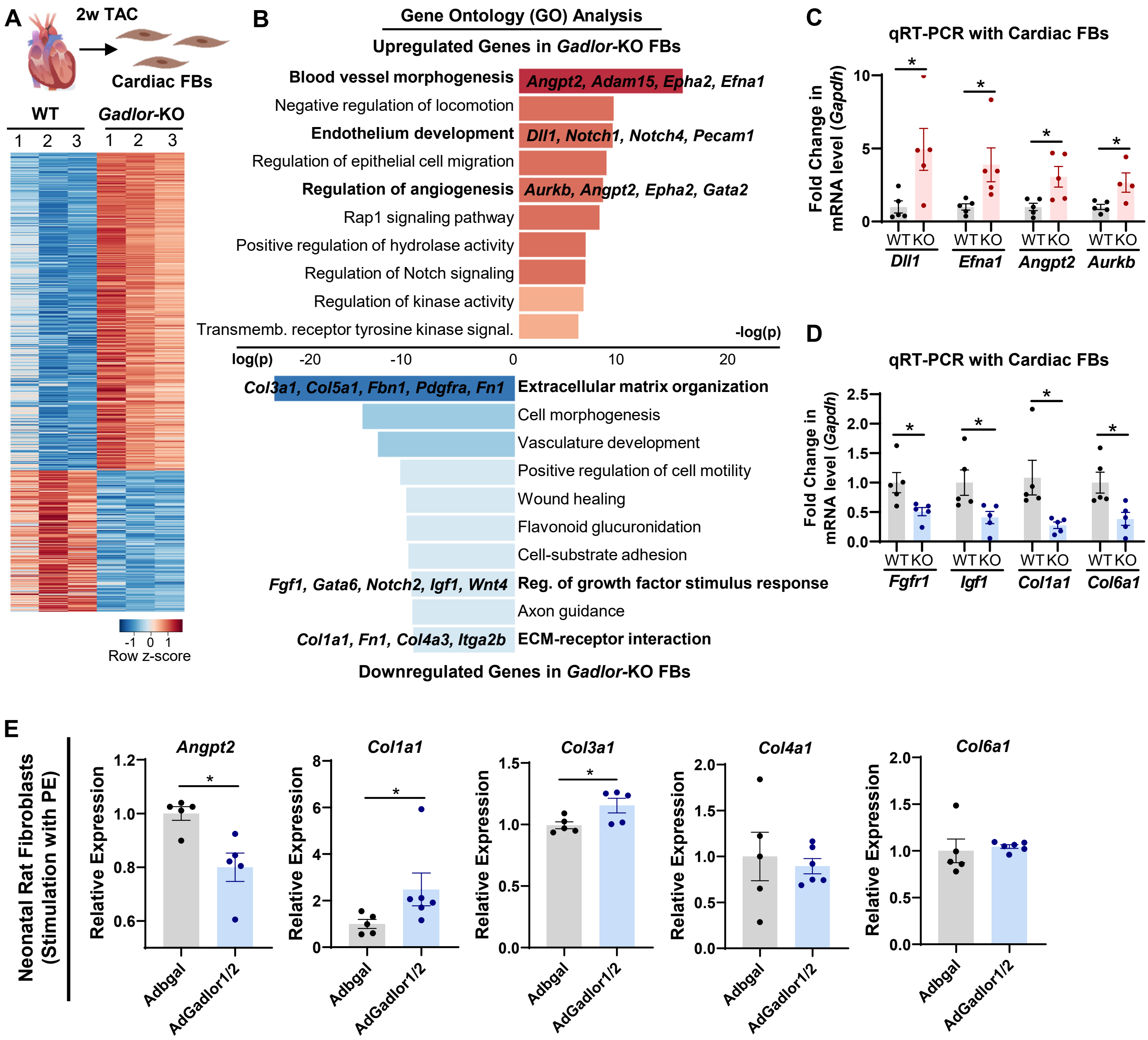


**Supplemental Figure 5: RNA sequencing of *Gadlor*-KO cardiac FBs showed less induction of fibrosis-associated genes after TAC.**

**A.** Heatmap showing differentially regulated genes analysed by bulk RNAseq of isolated cardiac fibroblasts after 2-weeks TAC. **B.** Bar plots showing the gene-ontology (GO) analysis of upregulated and downregulated genes in *Gadlor*-KO FBs after 2-weeks TAC (Red: Upregulated in *Gadlor*-KO, Blue: Downregulated in *Gadlor*-KO). Exemplary genes were listed for selected GO-terms. **C-D.** Validation of selected genes from RNAseq data with qRT-PCR in isolated adult cardiac FBs after 2 weeks of TAC (Red: Upregulated in *Gadlor*-KO, Blue: Downregulated in *Gadlor*-KO). **E.** qRT-PCR of selected genes in neonatal rat fibroblasts (NRFB) overexpressing *Gadlor* lncRNAs after *Gadlor1/2* adenovirus treatment (*βgal* as control) followed by phenylephrine (PE) stimulation (100 µM, 24 hours). Data are shown as mean±SEM. Data normality was evaluated with Shapiro-Wilk test and p-values were calculated with Student's t-test for parametric and Mann-Whitney test for non-parametric assessment. *p-value<0.05.

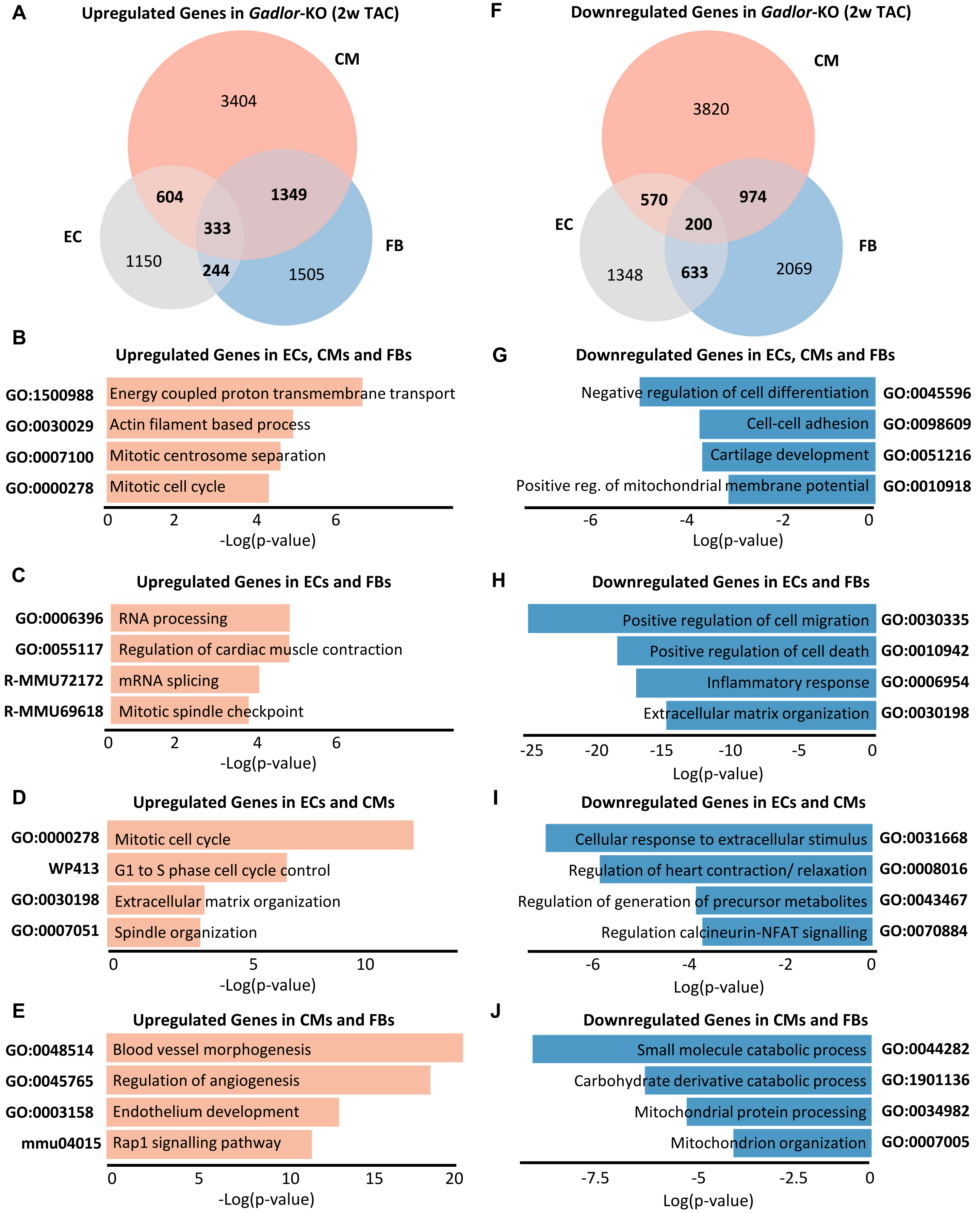


**Supplemental Figure 6: Venn diagrams of differentially expressed genes in different cardiac cell types of *Gadlor*-KO mice compared to WT littermates after 2 weeks of TAC. A.** Venn diagram of upregulated genes in different cardiac cell types (CM: red, EC: grey, FB: blue) of *Gadlor*-KO mice compared to WT after 2-weeks TAC. The numbers indicating the number of genes in corresponding cell types. Genes were filtered as fold-change greater or equals to 1.2 for each condition. **B-E.** Gene ontology (GO) analysis of indicated intersections representing the overlapping upregulated genes in shown cell type of *Gadlor*-KO compared to WT littermates. **F.** Venn diagram of downregulated genes in different cardiac cell types (CM: red, EC: grey, FB: blue) of *Gadlor*-KO mice compared to WT after 2-weeks TAC. **G-J.** Gene ontology (GO) analysis of indicated intersections representing the overlapping downregulated genes in shown cell type of *Gadlor*-KO compared to WT littermates. Bar plots showing selected GO-terms for each condition.

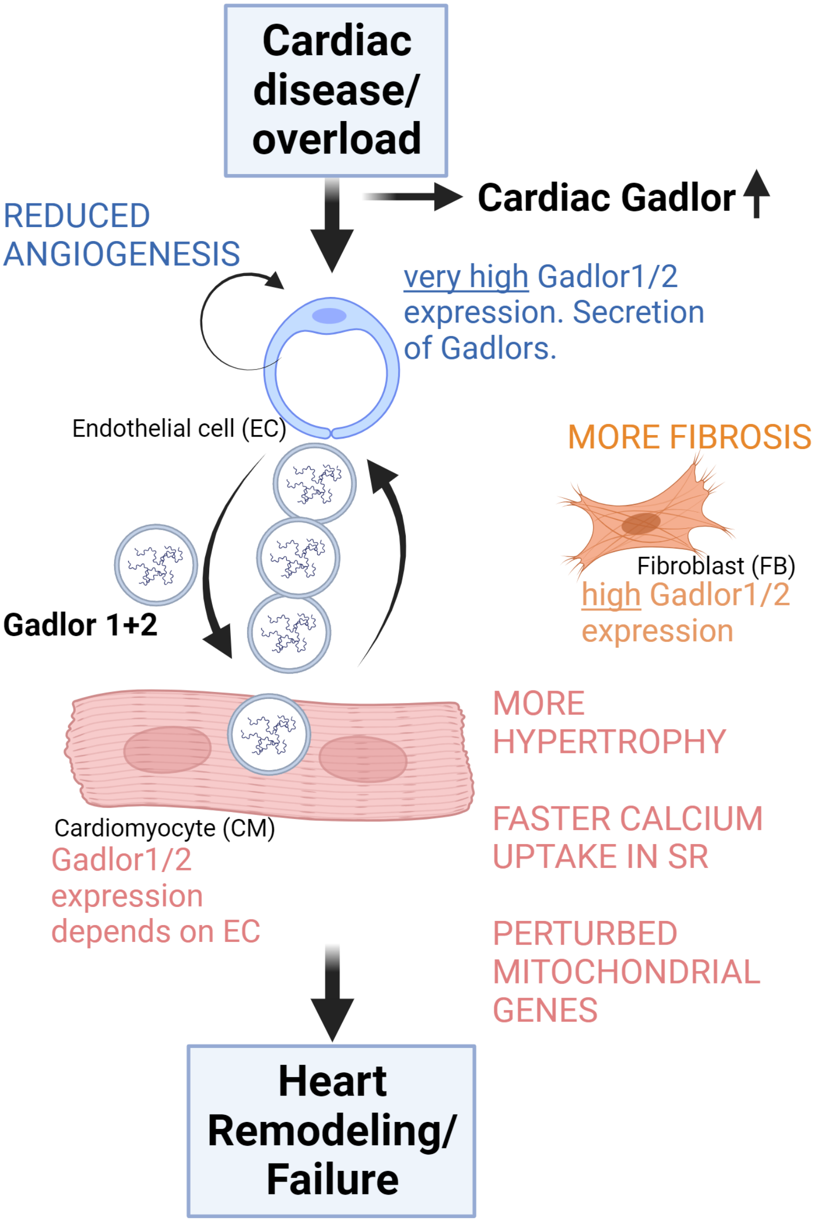


**Supplemental Figure 7:** Graphical summary showing *Gadlor1* and *Gadlor2* lncRNA expression and their effects (capitalized letters).The illustration was created with BioRender.com.

### Supplemental Tables

**Table 1:** Clinical information of human serum samples from patient cohort.

| Patient No | Age | Sex | Diagnosis | Previous Cardiac Surgery | NYHA Classification* During Admission | LVEF (%) |
|---|---|---|---|---|---|---|
| 1 | 77 | Female | AS, Hypertension, Dyslipidaemia, COPD | No | 3 | 40 |
| 2 | 73 | Male | AS, Hypertension, CAD, COPD | No | 3 | 40 |
| 3 | 90 | Female | AS, Hypertension, CAD | No | 3 | 45 |
| 4 | 86 | Female | AS, Hypertension | No | 2 | 55 |
| 5 | 80 | Male | AS, CAD | No | 3 | 35 |
| 6 | 91 | Female | AS, Hypertension, Dyslipidaemia, CAD, PAD | No | 3 | 60 |
| 7 | 75 | Male | AS, Hypertension, Diabetes, Dyslipidaemia | No | 3 | 50 |
| 8 | 86 | Female | AS, Hypertension, COPD | No | 3 | 45 |
| 9 | 83 | Female | AS, Hypertension, Dyslipidaemia, CAD | No | 3 | 65 |
| 10 | 76 | Male | AS, CAD | No | 4 | 30 |
| 11 | 84 | Male | AS, Hypertension, Diabetes, Dyslipidaemia, CAD, COPD | Yes | 3 | 40 |
| 12 | 85 | Male | AS, Hypertension | No | 3 | 20 |
| 13 | 80 | Male | AS, Dyslipidaemia, CAD | No | 3 | 25 |
| 14 | 76 | Male | AS, Dyslipidaemia, CAD | No | 3 | 45 |
| 15 | 76 | Female | AS, Hypertension, Diabetes, Dyslipidaemia, CAD | No | 3 | 40 |
| 16 | 82 | Female | AS, Hypertension, Diabetes, CAD | No | 3 | 65 |

*New York Heart Association (NYHA) Functional Classification
Criteria Committee, New York Heart Association, Inc. Diseases of the Heart and Blood Vessels. Nomenclature and Criteria for diagnosis, 6th edition Boston, Little, Brown and Co. 1964, p 114.
AS: Aortic Stenosis, CAD: Coronary Artery Disease, COPD: Chronic Obstructive Pulmonary Disease, LVEF: Left ventricle ejection fraction, PAD: Peripheral Arterial Disease.

**Table 2:** List of primers used for genotyping of *Gadlor*-KO mouse line

| | Primer 1 (Forward) | Primer 2 (Reverse) |
|---|---|---|
| ***Gadlor*-WT** | CTTGAGCCGTCTCTCCAAAG | GGGTGGCATGCAAGATGATTGAGA |
| ***Gadlor*-KO** | CTTGAGCCGTCTCTCCAAAG | TGTGGAGTGGACACATAGAGG |

**Table 3:** List of antibodies/reagents used in cell isolation methodologies and immunofluorescence (IF) staining.

| Target | Company, Catalogue No | Application |
|---|---|---|
| **Anti-mouse CD146 (LSEC) MicroBeads** | Miltenyi-Biotec, 130-092-007 | Cell isolation |
| **Feeder Removal MicroBeads, Mouse** | Miltenyi-Biotec, 130-095-531 | Cell isolation |
| **Isolectin B4 (IB4)** | Vector Lab, FL-1201 | 1:50 (IF) |
| **Purified Rat Anti-mouse CD102** | BD Pharmingen, 553326 | Cell isolation |
| **Purified Rat Anti-mouse CD31** | BD Pharmingen, 553370 | Cell isolation |
| **Rabbit Polyclonal anti-ki67** | Abcam, ab15580 | 1:100 (IF) |
| **VECTASHIELD HardSet Antifade Mounting Medium With DAPI** | Vector Lab, H-1500 | |
| **Wheat Germ Agglutinin (WGA)** | Invitrogen, W21405 | 1:100 (IF) |

**Table 4:** List of primers used for qPCR and qRT-PCR (5´-3´).

| | Primer 1 (Forward) | Primer 2 (Reverse) |
|---|---|---|
| *Gadlor1* | AGGTGAGCTCTGGTTGTGTT | CTGCTGCCTGTGAAAGATGG |
| *Gadlor2* | TGAGACTCCACTTGCCACAT | TGTGGTTTCAGGCATGTTTCT |
| *GADLOR1* | AATTTCAGCCACAAGCATCC | TGCTTGGGGAAGAGGAAGTA |
| *GADLOR2* | TGGGATCTAAGCACTGACACC | GAGACAGACATTCGTTTGGTCA |
| 18S | GTAACCCGTTGAACCCCATT | CCATCCAATCGGTAGTAGCG |
| U6 | CTCGCTTCGGCAGCACA | AACGCTTCACGAATTTGCGT |
| *Gapdh* | CCGCATCTTCTTGTGCAGT | CATCACCTGGCCTACAGGAT |
| *Acat1* | GCAGGGAAGTTTGCCAGTGAGA | GAACACGGTCTTGAGCTTTGGC |
| *Actn2* | CACCTGGAGTTTGCCAAGAGAG | GCCTTGAACTGCTCATGTGCAG |
| *Adam8* | TGCCAACGTGACACTGGAGAAC | GCAGACACCTTAGCCAGTCCAA |
| *Angpt2* | AACTCGCTCCTTCAGAAGCAGC | TTCCGCACAGTCTCTGAAGGTG |
| *Angptl4* | CTGGACAGTGATTCAGAGACGC | GATGCTGTGCATCTTTTCCAGGC |
| *Aurkb* | CTTCTACGACCAGCAGAGGATC | GGCATCTGACAGTTCCTCCATG |
| *Cacna1c* | CGTTCTCATCCTGCTCAACACC | GAGCTTCAGGATCATCTCCACTG |
| *Camk2d* | GTGACACCTGAAGCCAAAGACC | CCTGTGCATCATGGAGGCAACA |
| *Cdk1* | CATGGACCTCAAGAAGTACCTGG | CAAGTCTCTGTGAAGAACTCGCC |
| *Col15a1* | ACACCCACAGTGACTCCCAAGA | TCCTCATTGCCCACGATGTCTC |
| *Col1a1* | CCGCTGGTCAAGATGGTC | CCTCGCTCTCCAGCCTTT |
| *Col3a1* | ATAAGCCCTGATGGTTCTCG | ATGCATGTTTCCCCAGTTTC |
| *Col4a1* | ATGGCTTGCCTGGAGAGATAGG | TGGTTGCCCTTTGAGTCCTGGA |
| *Col6a1* | GACACCTCTCAGTGTGCTCTGT | GCGATAAGCCTTGGCAGGAAATG |
| *Comp* | GTGCCCAACTTTGACCAGAGTG | ACAGGCATCACCCACAAAGTCG |
| *Cox5a* | GTCACACGAGACAGATGAGGAG | CCGTCTACATGCTCGCAATGCA |

| | | |
|---|---|---|
| ***Cxcl2*** | CATCCAGAGCTTGAGTGTGACG | GGCTTCAGGGTCAAGGCAAACT |
| ***Dll1*** | GCTGGAAGTAGATGAGTGTGCTC | CACAGACCTTGCCATAGAAGCC |
| ***Efna1*** | GCTGAAGGTGACTGTCAATGGC | CGGCACTGTAACCAATGCTGTG |
| ***Fgfr2*** | GTCTCCGAGTATGAGTTGCCAG | CCACTGCTTCAGCCATGACTAC |
| ***Fh1*** | GAACTCACACGCAGGATGCTGT | GGCGGCTTTTATTCTCACCATCG |
| ***Fn1*** | TGTGACAACTGCCGTAGACC | TGGGGTGTGGATTGACCTTG |
| ***Gata4*** | GCCTCTATCACAAGATGAACGGC | TACAGGCTCACCCTCGGCATTA |
| ***Icam5*** | ACCGATGCACAGCAGTCAATGG | ATGTTCTGGGCAGCCTACACTG |
| ***Igf1*** | GTGGATGCTCTTCAGTTCGTGTG | TCCAGTCTCCTCAGATCACAGC |
| ***Il6*** | CGGCCTTCCCTACTTCACAA | TCCAGTTTGGTAGCATCCATCA |
| ***Klf15*** | ACACCAAGAGCAGCCACCTCAA | GCCTTGACAACTCATCTGAGCG |
| ***Lsamp*** | GGAGTCGAAGAGCAACGAAG | AATCTCAAGGCCATTTGCAC |
| ***Mfn2*** | GTGGAATACGCCAGTGAGAAGC | CAACTTGCTGGCACAGATGAGC |
| ***Myh6*** | GAGTGGGAGTTTATCGACTTCG | CCTTGACATTGCGAGGCTTC |
| ***Myh7*** | ACTGTCAACACTAAGAGGGTCA | TTGGATGATTTGATCTTCCAGGG |
| ***Nppa*** | TTCCTCGTCTTGGCCTTTTG | CCTCATCTTCTACCGGCATC |
| ***Nppb*** | GTCCAGCAGAGACCTCAAAA | AGGCAGAGTCAGAAACTGGA |
| ***Rcan1.4*** | CTTGTGTGGCAAACGATGATG | TGGTGTCCTTGTCATATGTTCTG |
| ***Sirt1*** | GGAGCAGATTAGTAAGCGGCTTG | GTTACTGCCACAGGAACTAGAGG |
| ***Sucla2*** | GGTGTCTCTGTTCCCAAAGGCT | TTTCCTCTGCCGCCAGCCAAAA |
| ***Tlr9*** | GCTGTCAATGGCTCTCAGTTCC | CCTGCAACTGTGGTAGCTCACT |
| ***ND1*** | CTAGCAGAAACAAACCGGGC | CCGGCTGCGTATTCTACGTT |
| ***16S*** | CCGCAAGGGAAAGATGAAAGAC | TCGTTTGGTTTCGGGGTTTC |

**Table 5:** List of *Gadlor1* and *Gadlor2* probes used for RNA antisense purification (RAP).

| | **Probe Sequence** |
|---|---|
| ***Gadlor1* – Probe 1** | [Btn]AGGATTGTTAAATATGACTATGCTTGGTATAGTCACAAAACATGGGAGTAC |
| ***Gadlor1* – Probe 2** | [Btn]TATCATAATCTTTCTGTAGGCCATTACTTGTTTCATATTTTAAAGGGACAG<br>TCCACTCTAGGAATGTCAAGTGTCTGATCTCTGAAAACA |
| ***Gadlor1* – Probe 3** | [Btn]TTTTCTGAATAGTTGAAAATTCTAACTAAACACAGGAAGAACAGAGNCAC<br>AAGAATAAAGAAATTTAGATATATCCTAAATGTTTCCAGG |
| ***Gadlor1* – Probe 4** | [Btn]ATAAAAGAAGGCGAGGGGTGGCATGCAAGATGATTGAGAAAGCCCAGT<br>AGCCATTTTTGGGGTGGGGCAAAGGGAGTGGTCTGGGTAGGG |
| ***Gadlor2* – Probe 1** | [Btn]GCTGCTATTTTATTATCTCTTTGGTTCTGTTTTTCATTTGTATTAAAAATG |
| ***Gadlor2* – Probe 2** | [Btn]GTGATGGTGAAGATGAAATTGAGATGAATCATTTGAAGAACGATGTGCGTT<br>TTAGAAGAATCACTTTGTC |
